# Leishmania Hijacks microRNA Import-Export Machinery of Infected Macrophage and Survives by Cross-Communicating Host miR-146a to Subjugate HuR and miR-122 in Neighbouring cells

**DOI:** 10.1101/2021.09.06.459146

**Authors:** Satarupa Ganguly, Bartika Ghoshal, Ishani Banerji, Shreya Bhattacharjee, Sreemoyee Chakraborty, Avijit Goswami, Kamalika Mukherjee, Suvendra N. Bhattacharyya

## Abstract

*Leishmania donovani*, the causative agent of visceral leishmaniasis, infects and resides within tissue macrophage cells of the mammalian host. It is not clear how the parasite infected cells cross-talk with the non-infected cells in the infection niche to regulate the infection process. Interestingly, miRNAs, the regulatory small RNAs of the host, could get trafficked into and out of infected cells as part of extracellular vesicles to ensure exchange of the epigenetic signals and can regulate the expression of their target genes in both donor and recipient cells. *Leishmania*, for its survival in host macrophage, adopts a dual strategy to regulate the intercellular transport of host miRNAs. The parasite, by preventing mitochondrial function of the host cells, restricts the entry of liver cell derived miR-122 containing extracellular vesicles in infected macrophage to curtail the inflammatory response by miR-122. The parasite reciprocally upregulates the extracellular export of anti-inflammatory miR-146a from the infected cells. The exported miR-146a restricts miR-122 production in liver cells and polarizes neighbouring naïve macrophages to the M2 state. miR-146a upregulates IL-10 in neighbouring macrophages where miR-146a dominates the RNA binding and miRNA suppressor protein HuR to inhibit the expression of proinflammatory cytokine mRNAs having HuR-interacting AU-rich elements and polarized the recipient cells to M2 stage.

**Graphical Abstract:** 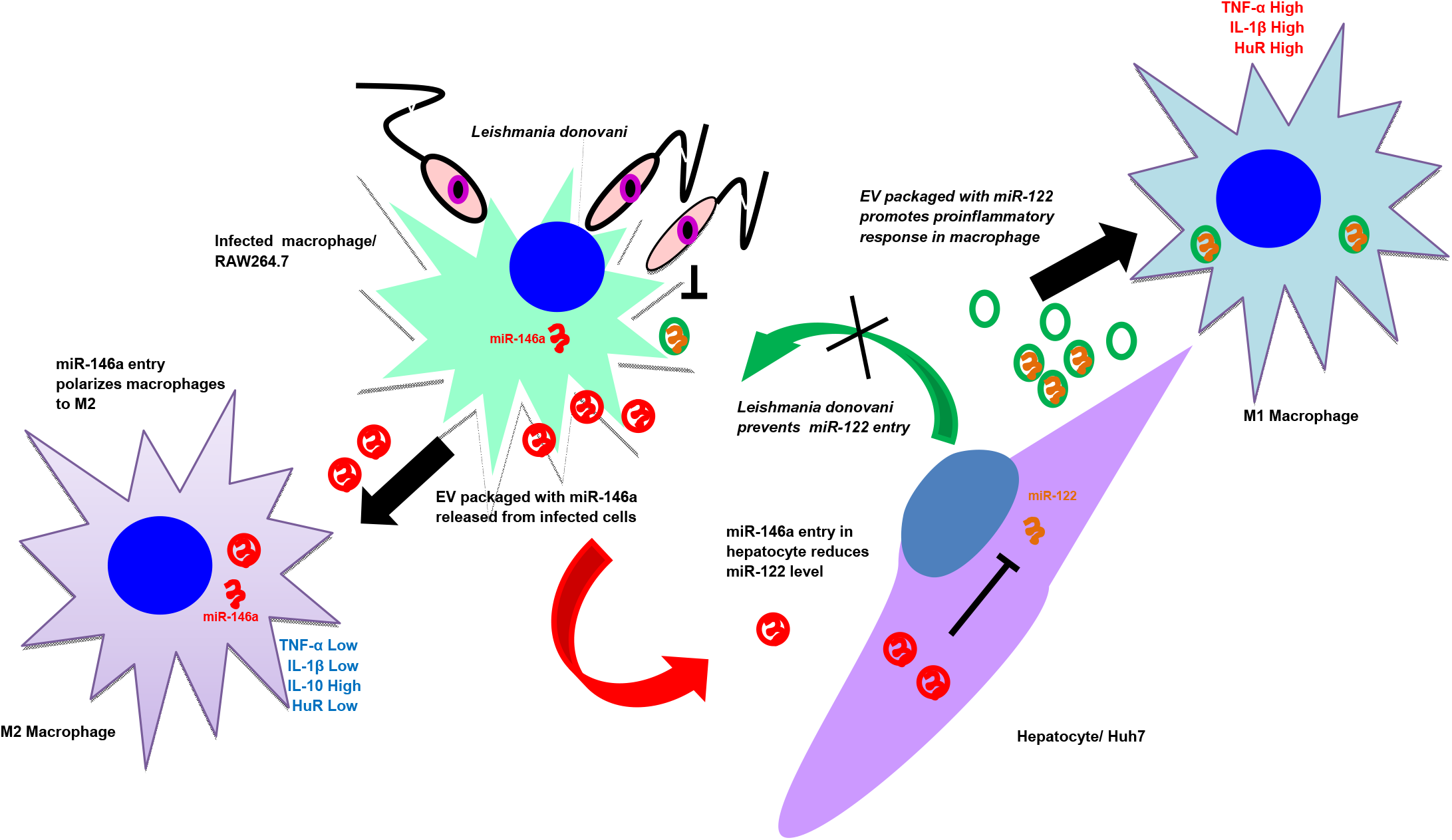

**Key Findings:**

- *Leishmania* stops vesicular entry of inflammatory miR-122 in infected cells by causing mitochondrial depolarization
- *Leishmania* secrets miR-146a from infected cells to stop miR-122 production in neighbouring hepatocytes
- miR-146a containing vesicles secreted by infected cells stops inflammatory response in recipient naïve macrophage
- miR-146 targets RNA binding protein HuR to stop inflammatory cytokine production

## Introduction

*Leishmania donovani* (*Ld*) is the causative agent of visceral leishmaniasis that affect a large portion of the human population in the Indian subcontinent and also in sub-Saharan Africa (Lenk et al, 2018). The apicomplexan parasite that has a dual stage life cycle lives in the gut of the sandfly vector as promastigotes. The promastigotes change to amastigotes within the mammalian host macrophage cells (Sunter & Gull, 2017). The Promastigotes enter the mammalian host with the saliva of the sand fly vector introduced during the blood meal, and subsequently make entry into the hepatic tissue where it infects the Kupffer cells at initial stages of infection before the infection load is transferred to the spleen (Walker et al, 2014). The *Ld* parasite lives in the tissue macrophages within a specialized sub-cellular structure called parasitophorous vacuole (Young & Kima, 2019)and alters the signalling components of the infected cells to polarize the infected macrophage to M2 stage. Thus *Ld* ensures low expression of pro-inflammatory cytokines like IL-1β and TNF-α while the anti-inflammatory cytokines IL-10 and IL-4 expression get enhanced (Mukherjee et al, 2013). The status of the non-infected macrophages present in the infection niche is not clear but it is certain that the non-infected macrophages present in the infection niche should not be activated to ensure the overall anti-inflammatory response being maintained in the infected tissue. How the parasite that remains within the specialized vacuole structure of the infected host macrophage cross-communicates with the resident non-infected macrophages to ensure suppression of cytokine expressionis an important question to explore. In this context, the extracellular signals derived from resident non-infected macrophages and hepatocytes that can activate the infected macrophage, should be counteracted by the infected host.

Extracellular Vesicles (EVs) are released by different types of mammalian cells that are used primarily by mammalian immune cells to cross-communicate the cellular status across the cell boundaries (Fernandez-Messina et al, 2015; Regev-Rudzki et al, 2013). It is known that the infected cell EVs can be used to transfer the parasite derived and host factors to target specific pathways in neighbouring cells to ensure establishment of systematic infection and propagation (Kalluri & LeBleu, 2020; Regev-Rudzki et al, 2013). miRNAs are important post-transcriptional regulators of gene expressions in mammalian hosts that are expressed in a cell type and stage specific manner (Bartel, 2018). miRNAs are also known to get communicated across the cell boundary to affect neighbouring cell fates. Therefore, if communicated from the infected host cells, miRNAs, as an epigenetic signal, could alter the gene expression in neighbouring cells. The role of specific miRNAs in regulation of expression of pro or anti-inflammatory pathways in mammalian immune cells have also been studied (Lindsay, 2008).

In this manuscript, we have documented an extraordinary mechanism that the internalized *Ld* adopted to ensure an anti-inflammatory infection milieu in the infected liver of the mammalian host. The internalized parasite prevents the pro-inflammatory response in the infected macrophage by preventing the entry of hepatocyte derived miR-122 containing EVs. The parasite achieves it through the depolarization of mitochondria of the host cells by enhancing the expression of the uncoupler protein Ucp2 that restricts the entry of the hepatic EVs into the infected cells and prevents miR-122 induced inflammatory responses. We have also noted miR-146a upregulation in the infected cells and the excess miR-146a also gets packaged into the EVs released by the infected macrophages. EVs with miR-146a enter the liver cells to restrict the production of miR-122. The miR-146a containing EVs are also internalized by the non-infected macrophages that then gets polarized to the M2 stage to express IL-10 in a miR-146a and HuR dependent manner. HuR, the RNA binding protein of known miRNA-derepessor function, induces export ofmiR-21 out of macrophage cells to cause high expression of miR-21 target IL-10 in the non-infected but miR-146a enriched cells. HuR, in turn itself, gets repressed by miR-146a. This in turn reduces the HuR-mediated stabilization and enhanced expression of pro-inflammatory cytokines in the recipient neighbouring macrophage cells. Thus, by cross communicating the infected host cell derived miR-146a, *Leishmania* ensures its own survival by regulating both miR-122 and HuR in neighbouring hepatocyte and naïve macrophage cells respectively.

## Results

### Secretion of miR-122 by activated hepatocytes

Lipopolysaccharide or LPS is an immunogen, derived from the cell wall of the gram-negative bacteria and is known to stimulate the macrophage cells via activation of TLR4 receptor and p38/MAPK pathway (Bode et al, 2012). LPS increases the expression of pro-inflammatory cytokines by enhancing the NF-ĸβ dependent transcription and also by inactivating the repressive miRNA pathway in LPS-activated macrophages (Mazumder et al, 2013). In mammalian liver, the LPS stimulation leads to activation of the tissue resident macrophages and thus, LPS increases liver inflammation (Rex et al, 2019). Hepatocytes also respond to LPS and altered metabolic function of hepatocytes exposed to LPS has been documented (Masaki et al, 2004; Momen-Heravi et al, 2015). miR-122 is the key miRNA expressed in hepatocytes, with known pro-inflammatory roles (Momen-Heravi et al, 2015). We documented a decreased miR-122 levels in mammalian liver cell Huh7 exposed to LPS (Supplementary Figure S1A-B). The decreased miR-122 level was associated with enhanced phospho-p38 level in LPS treated Huh7 cells (Supplementary Figure S1C). The decreased cellular miR-122 was associated with increased miR-122 detected in the EVs released by LPS exposed Huh7 cells (Supplementary Figure S1D-E). What consequence could this EV-associated miR-122 have on the tissue resident macrophages infected with *Leishmania* Upon infection of the host, the liver is the first tissue where the *Ld* initiates the infection process by targeting the tissue macrophage- the Kupffer cells (Beattie et al, 2010). miR-122 from hepatic cells, secreted as part of extracellular vesicles, can interact with macrophages to transfer the miRNA to the resident macrophages and could enhance the expression of pro-inflammatory cytokines there (Momen-Heravi et al, 2015). Therefore, the infected macrophages must adopt strategies to combat this activation process to protect themselves from getting killed by the immunostimulatory effect of hepatocyte secreted miR-122.

### *Leishmania* prevent internalization of proinflammatory miR-122 to the infected cells

*Ld* is also known to affect miRNA machineries of host cells to ensure its proliferation (Chakrabarty & Bhattacharyya, 2017; Ghosh et al, 2013). Interestingly, expression of miR-122 caused an increase in the TNF-α level in RAW264.7 macrophage cells (Figure 1A-C). Conversely, when *Ld* infection of RAW264.7 cells expressing miR-122 was followed, we found increased pro-inflammatory cytokine level and low anti-inflammatory cytokine levels upon miR-122 expression there (Figure 1D, E). The infection level of respective cells was also found to be reduced when the macrophages received miR-122 positive EVs before infection (Figure 1F, G). Thus, the hepatic miR-122 has an immuno-protective role and when transferred via EVs released by hepatic cells could prevent infection of neighbouring liver macrophage cells-the first target of invading *Ld* pathogen- in the mammalian host (Ghosh et al, 2013; Momen-Heravi et al, 2015). In order to survive, *Ld* must prevent this EV-mediated miRNA transfer process to stop inflammatory response in the host cells upon the interaction with miR-122 positive EVs. We hypothesized that *Ld* could have hijacked the inflammatory machinery of the host cell by preventing the miR-122 containing EV transfer to infected macrophages, and thus could ensure survival of the internalized pathogen. Consistent with the assumption, we found an increased production of pro-inflammatory cytokines and miR-155, the hallmark of inflammatory response in macrophage cells upon treatment with miR-122 positive EVs. But in cells already infected with *Ld*, the uptake of miR-122 containing EVs was found to be significantly compromised with concomitant reduction in miR-1555 or TNF-α production in the infection context (Figure 1H).

**Figure 1.**
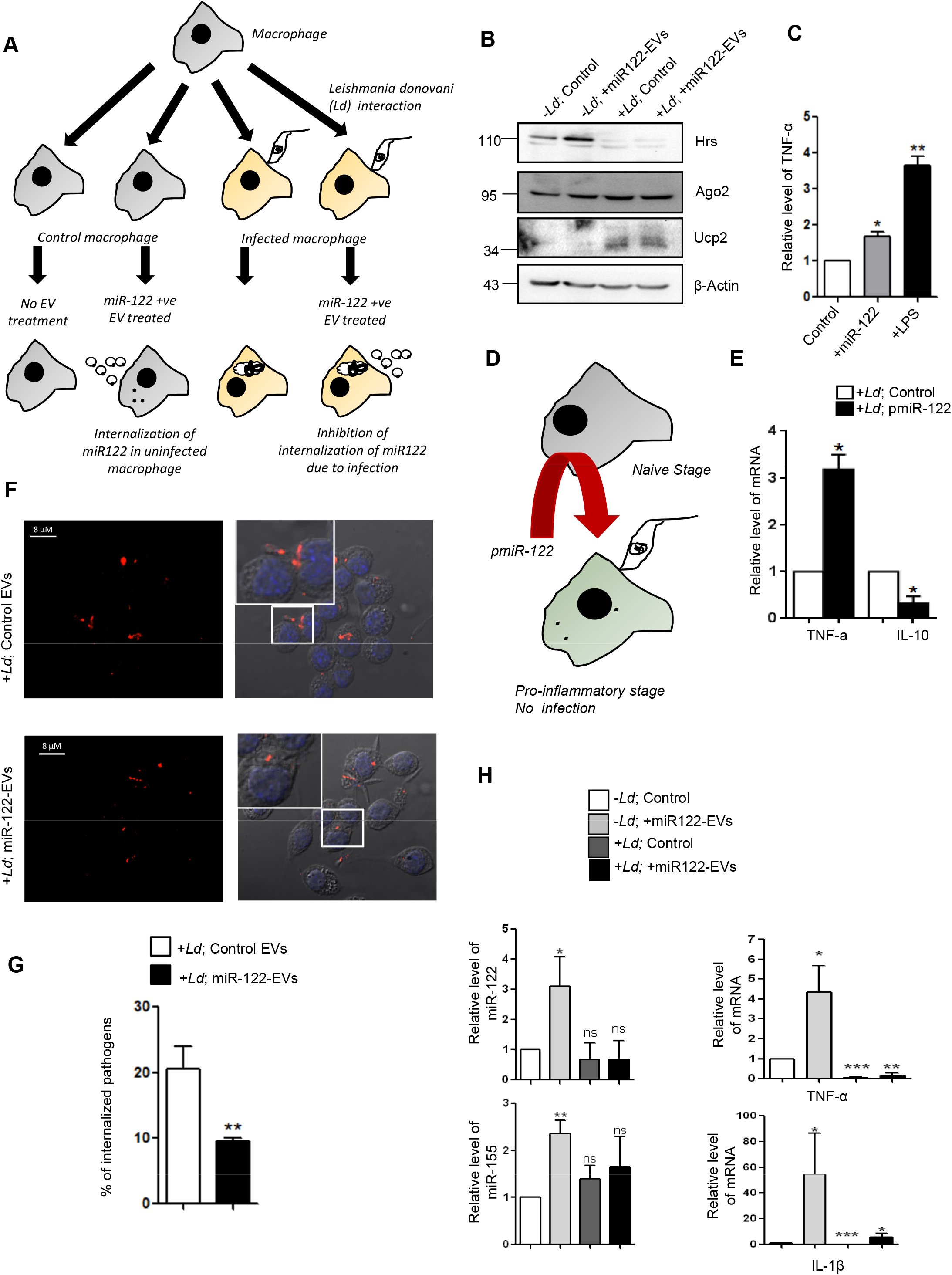
Hepatic miR-122 acts as immunostimulant for mammalian macrophage and *Ld* restricts miR-122 entry in infected macrophage. (A) Diagrammatic depiction of experiments done with *Leishmania donovani (Ld)* infected or uninfected RAW264.7 cells treated with miR-122 containing EVs. The macrophage cells were either infected or non-infected with *Ld* for 6 hours after which both the groups were either treated or untreated with miR-122 containing EVs. (B) Western blot analysis of RAW264.7 cells to show the infection related upregulation of Ucp2 and downregulation of HRS protein expression in RAW 264.7 cells (n=2). (C) Effect of miR-122 over-expression on cytokine expression in RAW 264.7 cells. The RAW264.7 macrophage cells were either kept as control, transfected with miR-122 expression plasmid pmiR-122 or treated with LPS (as positive control) followed by isolation of RNA to check the levels of pro-inflammatory cytokine, TNF-α. Quantification of TNF-α mRNA levels by qRT-PCR was done for the aforementioned conditions (n=3; p= 0.0361, 0.0084). (D) Effect of miR-122 expression on *Ld* infection. This panel shows the schematic model for RAW264.7 cells transfected with miR-122 expression plasmid followed by *L. donovani* infection. (E) Comparison of the proinflammatory TNF-α, and anti-inflammatory IL-10 cytokine expression in pmiR-122 transfected or pCI-neo control RAW264.7 cells followed by infection with *Ld*. An increase in the pro-inflammatory cytokines (p= 0.0180) and a decrease in the anti-inflammatory cytokine (p= 0.0422) was observed in the presence of miR-122 (n=3). (F) Microscopic analysis of *Ld* infection in RAW264.7 cells treated or untreated with miR-122 containing EVs. Cells were then visualised under the microscope. The Leishmania protein GP63 was labelled in red and cells with internalized parasites were counted. Scale bar 8 µm. Marked areas were zoomed 5x. (G) Effect of miR-122 containing EVs on *Ld* infection. Graphical representation of the percent of RAW264.7 cells infected with *Leishmania donovani* as observed microscopically. (n=132; p=0.0038). (H) Quantification of miRNA levels upon *Ld* infection followed by EV treatment in RAW264.7 cells. Real Time PCR showed no increase in miR-122 levels in infected macrophages incubated with miR-122 positive EVs (*top left panel;* p=0.0215). miR-155 levels were also quantified under similar conditions (*bottom left panel;* p=0.0024) (n=4). Relative levels of cytokine mRNA were also quantified. The proinflammatory cytokine, TNF-α (*top right panel;* p=0.0142) and IL-1β (*bottom right panel;* p=0.0438) did not increase in presence of the parasite followed by EV treatment (n=4). Error bars are represented as mean ± S.E.M. P-values were calculated by utilizing two-tailed Student’s t-test. ns: non-significant, *P < 0.05, **P < 0.01, ***P < 0.0001.

*Ld* is known to cause a robust change in endocytic pathway of the host cells and alters components that are also found to be used for endocytic entry or exit of miRNAs (Chakrabarty & Bhattacharyya, 2017; Lievin-Le Moal & Loiseau, 2016). Close proximity of endosomes with *Ld* containing parasitophorous vacuoles in infected macrophage cells was documented (Supplementary Figure S2A-B). We explored the status of Rab proteins in *Ld* infected macrophage and observe a decrease in endosomal protein HRS expression in infected cells (Figure 1B). However, there has not been a major change in Rab5a or RILP expression with *Ld* infection of RAW264.7 cells. Treatment of cells with EVs with miR-122 found to have no significant effect on endosome number or Rab5a expression in infected RAW264.7 cells. The infection also had no effect on Dynamin2 expression and thus cannot account for the reduced miR-122 entry observed in infected cells (Supplementary Figure S2B-C).

### *Leishmania* inhibits entry of miR-122 containing EVs in the infected Kupffer cells in mouse liver

EVs packed with miR-122 were generated from mouse liver cells HePa1-6. The miR-122 containing EVs were used to treat the mouse primary macrophage to score the effect of *Ld* infection on internalization of miRNA in infected mouse primary macrophages (Figure 2A). We documented a substantial reduction in EV-mediated miRNA entry in infected cells (Figures 2B). To score the same *in vivo*, we used mice infected with *Ld* and measured the effect of infection on EV-derived miR-122 entry in infected mouse liver macrophage cells (Figure 2C). After infection of 30 days and 24h of injection of miR-122 containing EVs, the hepatocytes and Kupffer cells were separated on a gradient and enrichment of each cell population was monitored by following the expression of mRNAs encoding the marker genes (Figure 2D). The levels of infection in both control and EV-injected population were monitored by following the kinetoplast DNA content (Figure 2E). In the uninfected Kupffer cells there was internalization of miR-122 after treatment with miR-122 containing EVs and a substantial increase in the expression of proinflammatory TNF-α mRNA in miR-122 EV-treated cells was noted (Figure 2F, G). However, the *Ld* infection substantially reduces the entry of miR-122 EVs in infected mouse Kupffer cells as both miR-122 and TNF-α expression was found to be substantially low in infected state (Figure 2H, I).

**Figure 2.**
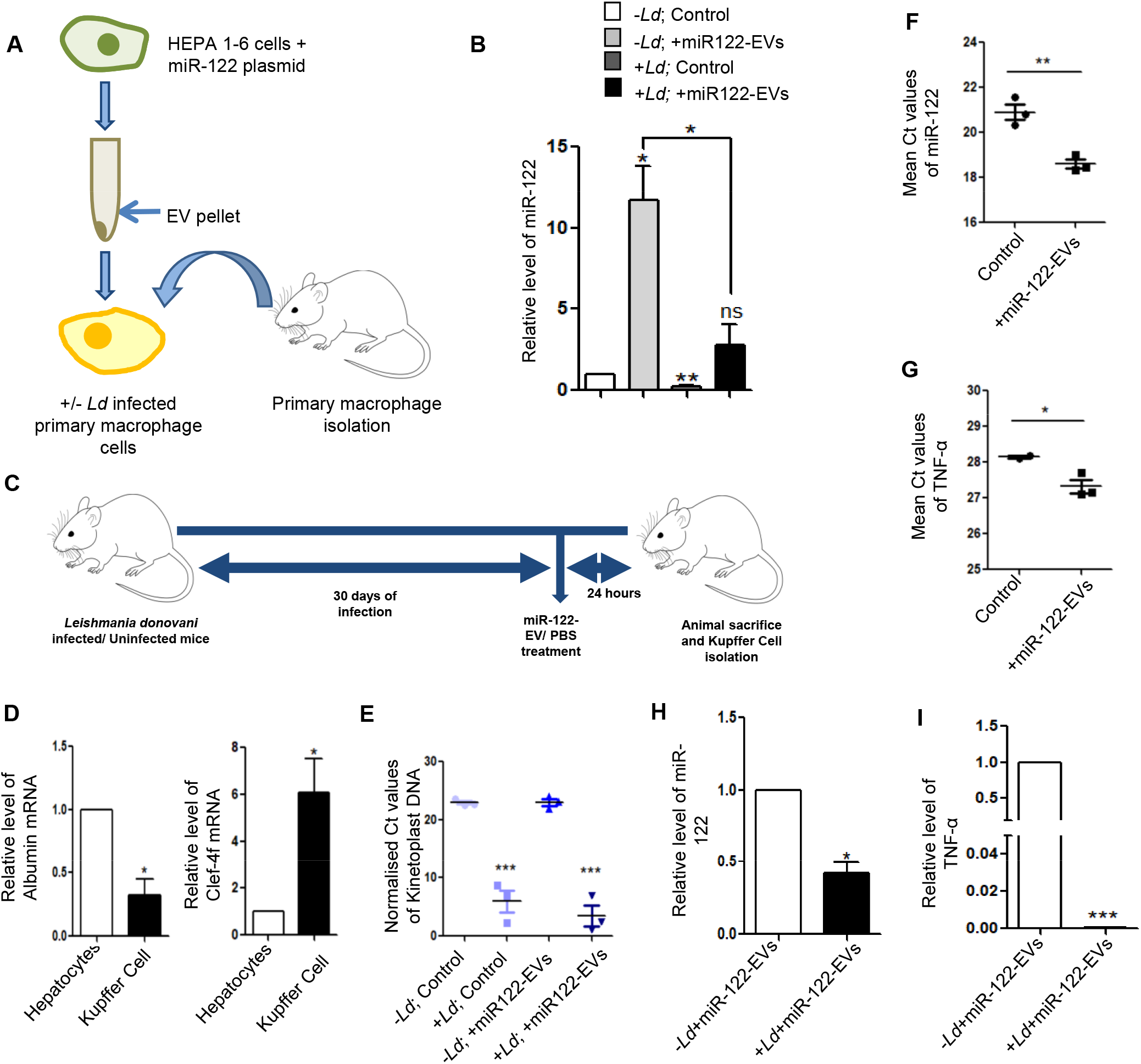
miRNA uptake is prevented in *Ld* infected primary macrophages and Kupffer cells. (A) A model to show the effect of *Leishmania* infection on uptake of miR-122-containing EVs in primary peritoneal macrophages. Macrophages were either infected or non-infected with *Ld* followed by miR-122-containing EV treatment. The EVs were obtained from miR-122 expressing murine hepatic cell Hepa 1-6. (B) Relative levels of internalized miR-122 in infected or non-infected primary macrophages (n=3; p=0.0379, 0.0227). (C) Schematic representation of *Ld* infection and treatment of mice followed by miR-122-containing EV treatment. Mice were infected with *Ld* for 30 days followed by tail vein injection of miR-122 positive EVs. After 24 hours of injection mice were sacrificed and Kupffer cells were isolated to quantify the miR-122 content. (D) Characterisation of isolated hepatocytes and Kupffer cells. Quantification of hepatocyte specific Albumin mRNA in hepatocytes and Kupffer cells to show their low levels in Kupffer cells (p= 0.143; *left panel*). Relative levels of macrophage specific C-type Lectin Domain Family 4, Member F (Clec4f) mRNA between hepatocytes and Kupffer cells showing high levels in Kupffer cells (p= 0.0378; *right panel*) (n=4). (E) Comparison of normalised Ct values of Kinetoplast DNA in infected or uninfected Kupffer cells and untreated or treated with EVs containing miR-122 to determine the parasitic load in Kupffer cells (n=3; p=0.0009, 0.0005). (F) Mean Ct values of internalized miR-122 in uninfected Kupffer cells either untreated or treated with miR-122 containing EVs (n=3; p=0.0049). (G) Mean Ct values of TNF-α mRNA in uninfected Kupffer cells upon treatment with miR-122-EVs compared to untreated Kupffer cells (n=2; p=0.0444). (H) The internalization of EV derived miR-122 declined in the presence of *Leishmania* infection. Real Time PCR revealed the relative levels of miR-122 in uninfected or infected Kupffer cells but treated with miR-122 containing EVs (n=3; p=0.0162). (I) Relative levels of pro-inflammatory cytokine TNF-α in *Leishmania* infected or uninfected but miR-122-EV treated Kupffer cells (n=3; p<0.0001). Error bars are represented as mean ± S.E.M. P-values were calculated by utilizing two-tailed Student’s t-test. ns: non-significant, *P < 0.05, **P < 0.01, ***P < 0.0001.

### *Leishmania* depolarizes mitochondria to prevent the entry of miRNA-containing EVs

*Ld* targets the mitochondrial dynamics and activity by inducing depolarization of mitochondria and reducing the mitochondria-ER-endosome interaction in infected cells (Chakrabarty & Bhattacharyya, 2017). Mitochondrial uncoupler protein Ucp2 gets upregulated upon *Ld* infection (Figure 1B) suggesting a depolarized mitochondrial state in infected cells (Chakrabarty & Bhattacharyya, 2017). Does mitochondrial depolarization affect EV-entry? Mitochondria can interact with several cellular compartments, like ER and endosomes (Klecker et al, 2014; Todkar et al, 2019). The Ucp2 is a mitochondrial uncoupling protein causing defects in oxidative phosphorylation and ATP synthesis by changing membrane potential across mitochondrial membranes. Hence, it was anticipated that uncoupling of the mitochondrial potential can affect the internalization of EV-derived miRNAs in mammalian cells as it is known to affect endogenous miRNA activity (Chakrabarty & Bhattacharyya, 2017). FA-Ucp2 was expressed in the recipient HeLa cells and FA-Ucp2 expressing cells were incubated with miR-122-containing EVs. Ucp2 overexpression caused a disruption in the mitochondrial structures as observed microscopically (Figure 3A). The low level of internalization and consequently a lower repression activity of the transferred EV derived miRNA were observed in FH-Ucp2 expressing recipient HeLa cells (Figure 3B-D). It has been reported earlier that *Ld* infection or the loss of mitochondrial membrane potential by FA-Ucp2 are accompanied by reduced juxtaposition of endoplasmic reticulum (ER) and mitochondria (Chakrabarty & Bhattacharyya, 2017)(Supplementary Figure S3A, S3B). Mfn2 is a protein responsible for mitochondrial tethering with ER. Mfn2 knockout mouse embryonic fibroblasts (MEFs) showed a decrease ER tethering of mitochondria. To revalidate the Ucp2 overexpression mediated loss of ER-mitochondria contact, being responsible for the reduced miRNA uptake, EV associated miRNA internalization and Ago2 association were compared in Mfn2 knockout and wild type MEFs. Importantly, as reported earlier, the endogenous miR-16 level was found to be increased in Mfn2 depleted cells (Figure 3E-F) that has been consistent with the previous findings on increased cellular miRNA content upon Ucp2 over-expression (Chakrabarty & Bhattacharyya, 2017).

**Figure 3.**
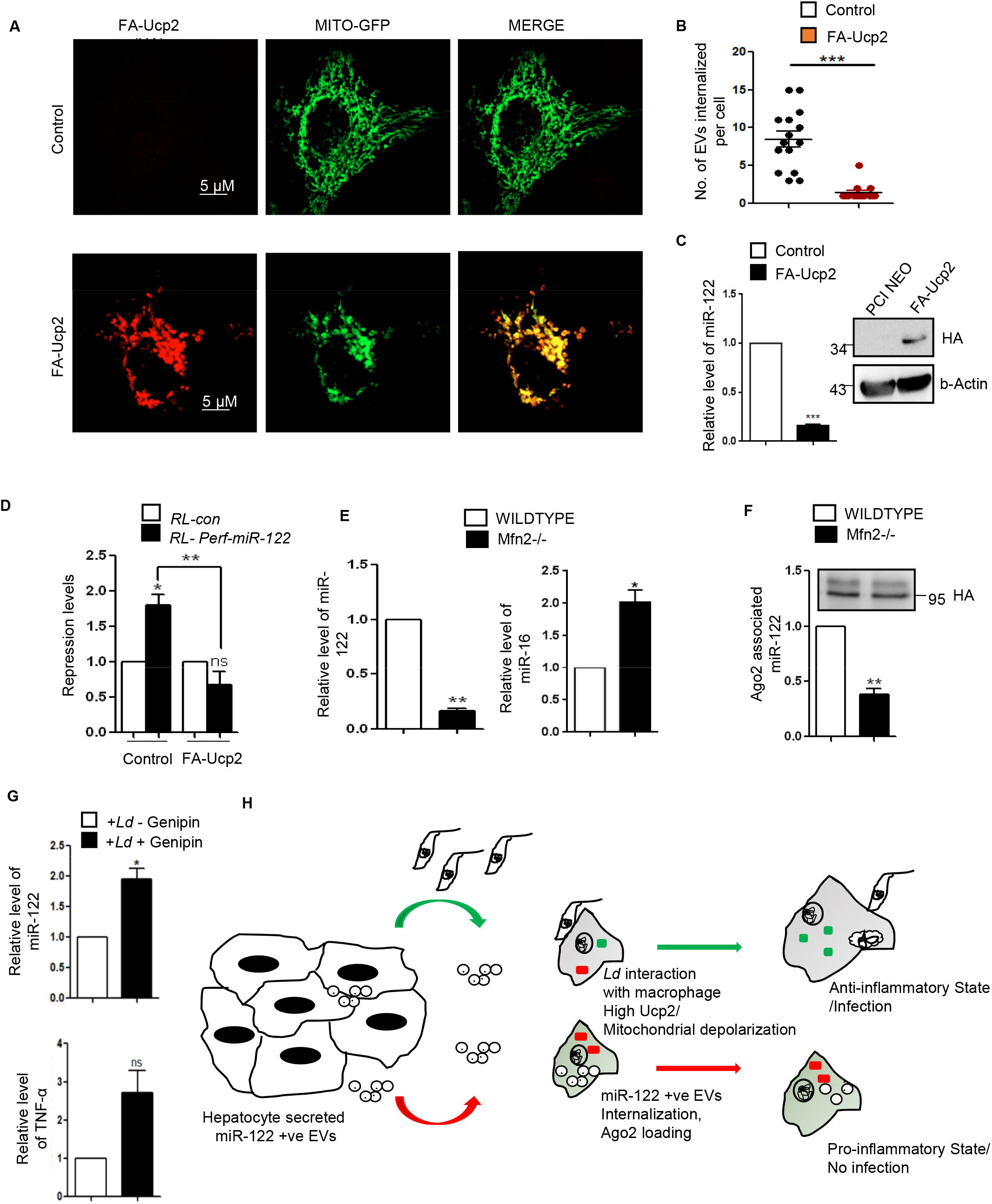
Mitochondrial depolarization targeted by *Ld* prevents internalization and activity of EV-derived miRNAs in recipient cells. (A) Structure of mitochondria (green) in Mito-GFP expressing HeLa cells transfected with FA-Ucp2 (red) or control plasmid. Scale bar 5µm. (B) Graphical representation of the internalization of CD63-GFP positive EVs in recipient HeLa cells transfected either with control or FA-Ucp2 expressing plasmids (n=18; p<0.0001). (C) Effect of Ucp2 overexpression on uptake of miR-122. Relative level of internalized EV derived miR-122 in recipient HeLa cells transfected either with control or FA-Ucp2 expressing plasmid (*left panel*; n=3; p<0.0001). *Right panel* shows the western blot for HA. (D) Luciferase assay to show the repression levels of RL-perf-miR-122 in recipient cells transfected with control plasmid or FA-Ucp2 and treated with miR-122 containing EVs. The repression levels were calculated as a ratio of firefly normalised RL-Control value to firefly normalised RL-perf-miR-122 value (n=3; p=0.0013). (E) Relative level of internalized miR-122 (*left panel;* p=0.0012) and endogenous miR-16 levels (*right panel;* p=0.0316) in Mfn2 (Mitofusin2) wildtype and knockout mouse embryonic fibroblast (MEF) cells treated with miR-122 containing EVs. Quantification was done by qRT-PCR and U6 levels were used as normalizing control (n=3). (F) FA-Ago2 associated miR-122 levels in Mfn2 wild type and knockout MEF cells (transfected with FA-Ago2) co-cultured with miR-122 expressing HeLa cells. Immunoprecipitated Ago2 levels were used for normalization of associated miR-122 (n=3; p=0.0077). (G) Rescue of EV-miR-122 internalization in *Leishmania donovani* infected RAW cells in presence of Genipin, the inhibitor of Ucp2. RAW cells infected with *L. donovani* were treated with Genipin (100µM after 4 hours of infection) followed by addition of miR-122 positive EVs. Relative levels of miR-122 (*top panel;* p=0.0311) and TNF-α mRNA levels (*bottom panel;* p=0.0968) increased in infected RAW264.7 cells treated with Genipin (n=3). (H) Diagrammatic representation of the EV-mediated crosstalk between hepatocytes and macrophages. Hepatocytes release miR-122 containing EVs which can be transferred to macrophages that causes production of proinflammatory cytokines and can prevent Ld infection (red arrow). Inversely, *Ld* infected macrophages are unable to take up the miR-122 containing EVs due to upregulation of Ucp2 protein in cells and are tuned to have high production of anti-inflammatory cytokines for sustained parasitic infection (green arrow). Error bars are represented as mean ± S.E.M. P-values were calculated by utilizing two-tailed Student’s t-test. ns: non-significant, *P < 0.05, **P < 0.01, ***P < 0.0001.

The miR-122 entry and associated TNF-a expression has been observed in infected macrophage cells treated with Ucp2 inhibitor Genipin (Figure 3G). Overall, this data suggest *Ld* infection associated increase of Ucp2 to cause mitochondrial depolarization to prevent the entry of the proinflammatory miR-122 entry via EVs into the infected macrophage to stop expression of inflammatory cytokines (Figure 3H).

### *Leishmania* infection increases cellular and extracellular miR-146a levels in infected cells

The hypothesis, that the infected cells should release the EVs enriched with factors to facilitate the propagation of infection, inspired us to examine the key protein components that are specifically released by infected cells or export of which is prevented in the infection context. The EVs isolated from control and *Ld* infected cells were analysed. Although upon infection there has been a drop in expression of HRS or Rab27a proteins that are known to have role in late endosome/MVB formation and EV release (Figure 4A-B), we did not detect a significant change in EV- diameter or marker protein levels in isolated population from both control and infected RAW264.7 cells (Figure 4C-E). In the mass spectrometric analysis followed by candidate protein identification, we have documented several candidate proteins that showed exclusive presence in the EVs isolated from control or *Ld* infected cells. However, differential expression of any important regulatory factors of cytokine expression were not detected (Supplementary Figure 4, Supplementary Table 1-3). Therefore, it could be the small RNA population of the infected host that may carry the required information to the neighbouring cells as part of EVs. In infected cells, both miR-146a and miR-155, two most important regulatory miRNAs of inflammation pathways, were found to be upregulated. Interestingly, among them, only miR-146a were detected in EVs released by infected cells and the content of miR-146a had increased several folds in EVs from *Ld* infected cells compared to EVs from control non-infected cells (Figure 4F-G). Similar increase of miR-146a in EVs released by infected primary macrophage was also detected (Figure 4H). The miRNAs that are reported to be increased in *Ld* infected cells (Chakrabarty & Bhattacharyya, 2017), miR-21, miR-125b or miR-16 all were found to be increased in the EVs released by infected cells (Figure 4I).

**Figure 4.**
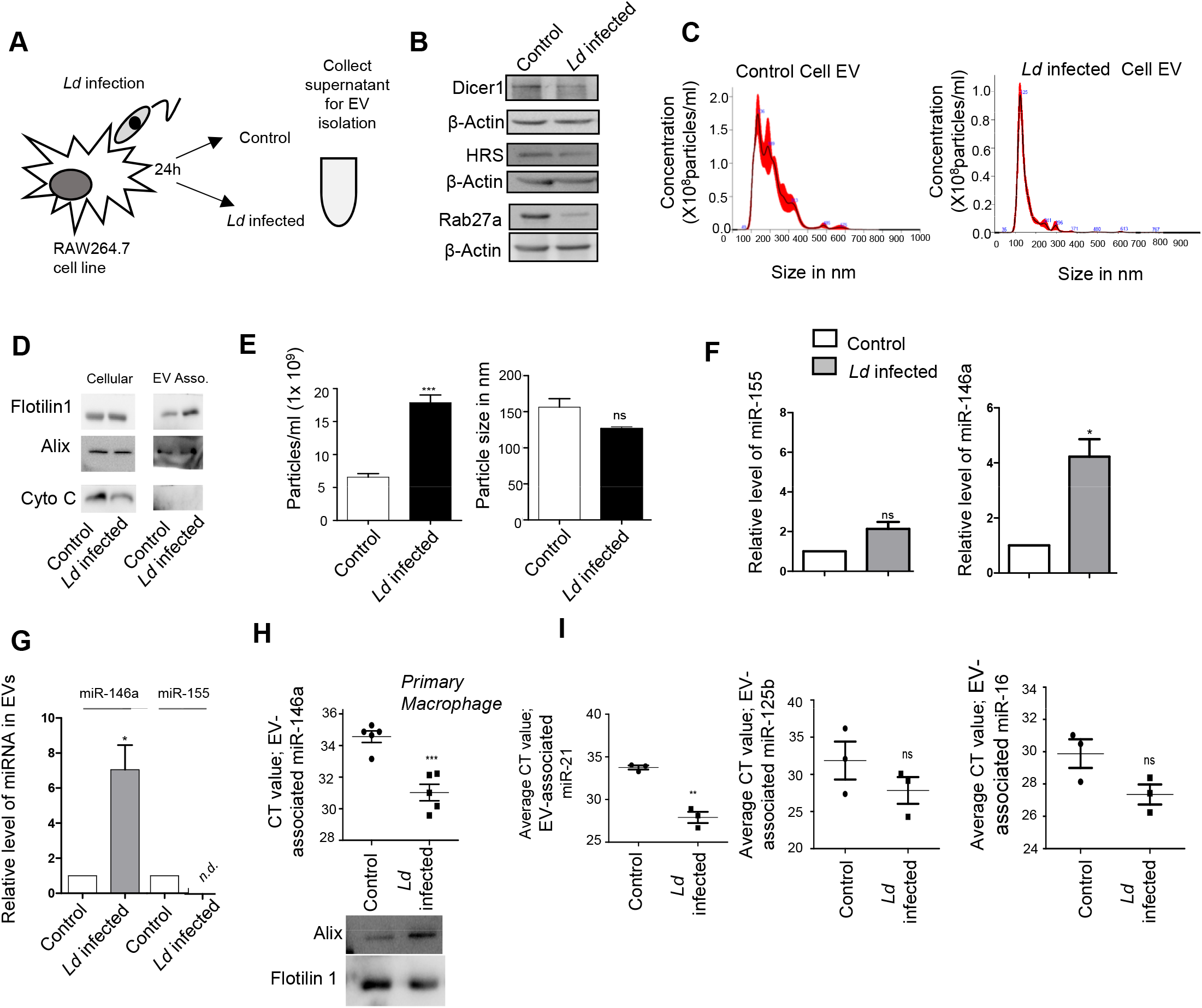
*Leishmania donovani (Ld)* infection triggers extracellular export of miRNAs from infected macrophage. (A) Schematic representation of the experiment. After 24h of *Ld* infection the host macrophage and EVs isolated from culture supernatant were analysed for cellular and EV associated protein and RNA analysis. (B) *Ld* infection altered cellular protein levels. Western blot images showed protein levels after 24 h of infection of macrophages. β-Actin was used as loading control. (C-E) Characterization and quantification of Extracellular Vesicles (EV) released from control and *Ld* infected macrophages. Nanoparticle Tracking Analysis (NTA) of EVs released from control (C, left panel) and infected cell (C, right panel). Western blot images showed EV marker proteins in control and infected cells and released EVs (D). Number and size of EVs released by control and infected cells were quantified by NTA analysis (E, left panel, p=0.0008, N=3 & right panel, p=0.0677, N=3, respectively, unpaired *t*-test) (mean+/− S.E.M). (F-G) miR-146a increases and is exported from *Ld* infected macrophage cells. qRT-PCR data represented the relative level of miR-146a and miR-155 in EVs from RAW264.7 cells. EV marker protein alix was used for normalization of EV associated miR-146a level (F, p=0.0229, N=4). qRT-PCR based relative quantification of cellular miR-155 (G, left panel, p=0.0817, N=3) and miR-146a (G, right panel, p=0.0146, N=4) after 24hour infection of RAW264.7 cells. U6 was used as endogenous control. miRNA levels of infected cells were normalized against non-infected controls. (H) *Ld* infection triggers the extracellular export of miRNAs from mouse peritoneal macrophages. miR-146a level was also validated in EV derived from mouse peritoneal macrophage (H, upper panel, p=0.0006, N=5, unpaired *t*-test) and western blot data also showed EV marker proteins (H, lower panel). (I) Relative change in amount of miR-21 (I, left panel, p=0.0011, N=3), miR-125b (I, middle panel, p=0.2654, N=3) and miR-16 (I, right panel, p=0.0794, N=3) in EVs released from *Ld* infected and non-infected RAW264.7 cells (I, unpaired *t*-test) (mean+/− S.E.M). In all experimental data, ns: non-significant, *, ** and *** represent *P*-value of <0.05, <0.01 and <0.0001 respectively calculated by student paired *t-*test where not mentioned otherwise.

### *Leishmania* by targeting HuR ensures export of miR-146a but not miR-155 from the infected cells

We report differential export of miR-146a from infected RAW264.7 macrophage. But how miR-146a gets differentially exported out of infected macrophage cells is a fascinating question. Retention of miRNA and their targets with polysomes has been found to be the reason for reduced miRNA export noted in mammalian cells (Ghosh et al, 2015) while target RNA presence positively influences the export process (Ghosh et al, 2021). We have isolated the polysomes from control and infected cells and detected increased retention of both miR-146a and miR-155 with polysomes in the infected cells (Supplementary Figure 5A, D). The increased retention of IL-10 and MyD88 mRNAs, known to get expressed in infected cells, were also detected more with polysomes in the infected cells but the miR-146a target TRAF6 mRNA and TNF-α mRNA was found to be less with infected cell polysomes (Supplementary Figure 5 B and C).

The polysome retention data thus could not clearly explain why there has been a preference of miR-146a for export from *Ld* infected cells. miRNAs accumulate at endosomes before they get packed and exported out (Mukherjee et al, 2016). We had isolated the endosomes and ER fractions from control and infected cells to document preferential enrichment of miR-146a over miR-155 at the endosomal fraction that can explain its enhanced export observed in infected macrophage cells (Supplementary Figure 5 E-G). HuR is known for its role in export of miRNAs in mammalian liver and macrophage cells. HuR drives miRNA accumulation with endosomes before the export but HuR level is known to get reduced in *Ld* infected cells (Goswami et al, 2020) (Supplementary Figure 5H). As HuR level increases with LPS treatment which is associated with enhanced export of miR-155 from LPS activated macrophage cells (Goswami et al, 2020) it could be possible that low HuR in *Ld*-infected cells could cause reduced export of miR-155. Could it be possible that binding of HuR with miR-155 makes it available in endosomal fraction and export? Supporting the notion, we have documented strong interaction of HuR with miR-155 in LPS treated cells (Supplementary Figure 5I-J). Similarly export of miR-155 from HuR expressing cells has also been noted (Supplementary Figure 5K).

### EV-containing miR-146a downregulates miR-122 maturation in hepatocytes

The infected cells prevent the miR-122 containing EVs entry and restricts the activation of the *Ld*-infected cells by miR-122 in order to prevent the death of the internalized parasites due to enhanced pro-inflammatory response observed in miR-122 recipient macrophage (Chen et al, 2011). However, miR-122 in hepatic EVs could also get transferred to a naïve neighbouring macrophage not infected with *Ld*. How the parasite could prevent the activation of resident neighbouring naïve macrophage also to prevent the pro-inflammatory cytokine accumulation in the infection microenvironment? It is likely that *Ld* must have adopted a mechanism to reduce the miR-122 containing EV release by neighbouring hepatocytes in the infection niche to prevent miR-122 mediated activation of naïve non-infected macrophage cells. Can *Ld* do it through cross talk with hepatocytes via the EVs released by infected cells? We have noted miR-146a as the predominant anti-inflammatory miRNA in the EVs secreted by the infected macrophage (Fu et al, 2017). Does miR-146a reduce the miR-122 level in the hepatocytes?

We have isolated the EVs from infected macrophage for treating human hepatoma cell Huh7. We have documented an increase in miR-146a content of recipient hepatic cells with a reduction in mature miR-122 level there (Figure 5A, B). The decrease of mature miR-122 was accompanied by an increase in pre-miR-122 level and with an increase in endogenous control miR-16 level in the hepatocyte. Thus, the reductive effect of miR-146a on hepatic miR-122 is a miRNA specific effect. To confirm that the effect of miR-146a on mature miR-122 level is miR-146a specific, we have isolated the EV from RAW264.7 cells ectopically expressing miR-146a and used the isolated EVs for treatment of the hepatocytes (Figure 5C). The EVs isolated from control and pmiR-146a expression plasmid transfected cells were analysed by NTA to document no major change in size and number of the EVs under miR-146a expression condition (Supplementary Figure S6A). Increased miR-146a content in EVs released from pmiR-146a expressing cells was also noted (Supplementary Figure S6B) and cellular and EV fractions were western blotted for EVs and cytosolic markers to confirm the purity of the fractions (Supplementary Figure S6C). The miR-146a containing EV treatment enhanced cellular miR-146a level in recipient hepatocytes with concomitant decrease of mature miR-122 and increase of pre-miR-122 levels (Figure 5D). To substantiate the idea of miR-146a induced reduction in miR-122 levels in hepatocytes, we have expressed pmiR-146a in hepatocytes and documented decreased miR-122 level associated with increased pre-miR-122 and miR-146a levels (Figure 5E). We have noted a reduction in Dicer1 level upon *Ld-*infected cell-EV treatment of hepatocytes. However, miR-146a-containing EV-treatment or miR-146a expression in hepatocytes do not alter Dicer1 level (Figure 5F-H). Thus, altered pre-miR-122 processing by Dicer1 in miR146a enriched hepatocyte could not be explained by the unhanged Dicer1 level. The increased pre-miR-122 however get associated with the Dicer1 to a lesser extent in cells expressing pre-miR-146a (Figure 5J and K). Therefore, there must be a mechanism that excludes the pre-miR-122 to be associated with Dicer1 for its processing to the mature form. The miR-146a expression however enhances the p-ERK level and a decrease in p-p38 level (Figure 5I), suggesting a possible lowering of transcriptional events for pre-miR-122.

**Figure 5.**
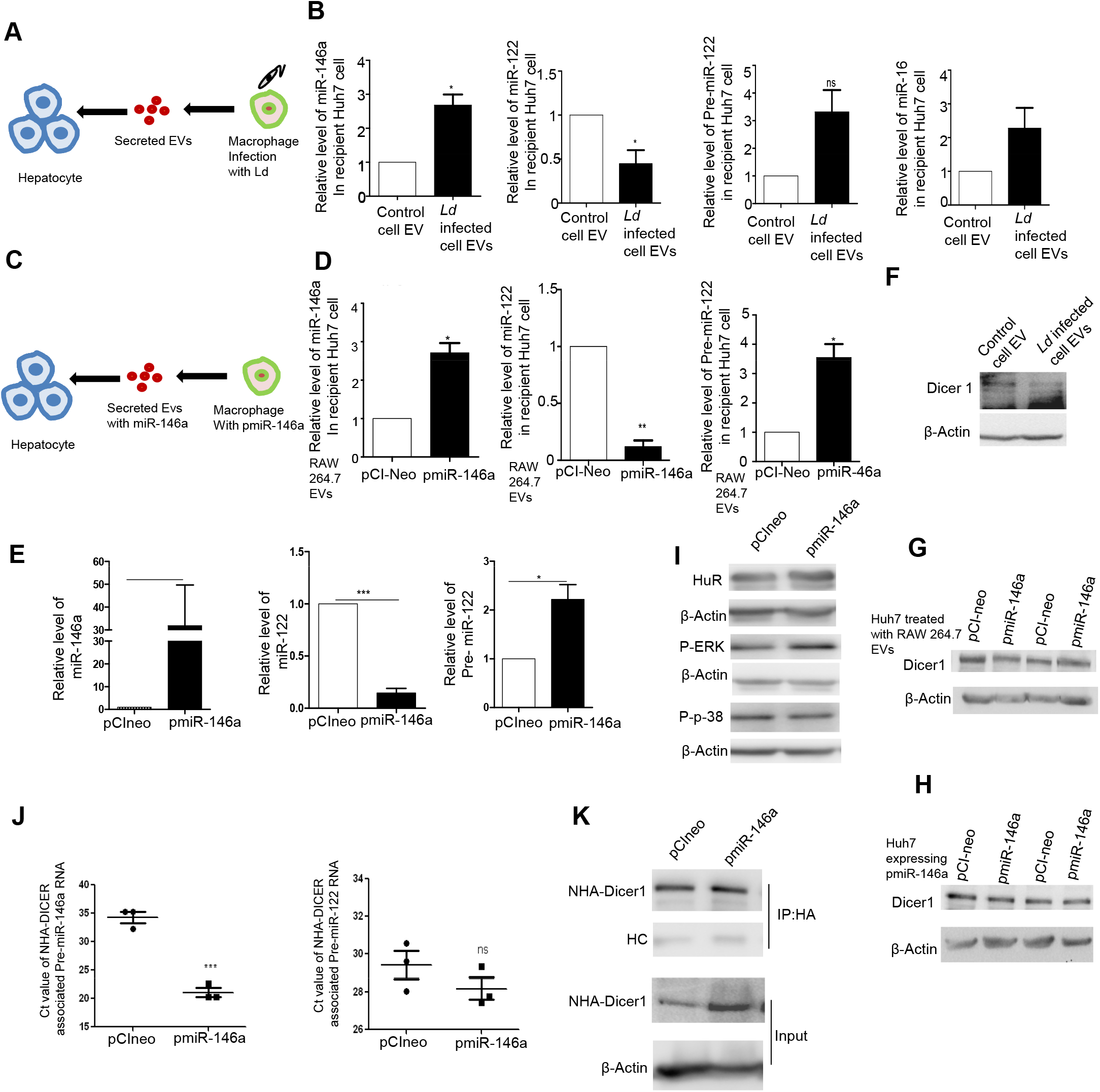
EV mediated transfer of miR-146a reduces miR-122 level in hepatic cells. (A-B) Effect of control and Ld-infected cell derived EVs on naïve hepatocytes. A schematic representation of the experiment (A). Relative levels of miR-146a (B, left panel, p=0.0329, N=3), miR-122 (B, middle left panel, p=0.0364, N=4), Pre-miR-122 (B, middle right panel, p=0.0993, N=3), miR-16 (B, right panel, p=0.2796, N=2) was measured by qRT-PCR in recipient hepatic Huh7 cells after 24 hours of control and infected cell derived-EV treatment. U6 was used as endogenous control for miRNA normalization and miRNA levels of control EV-treated cells was used for normalization of infected cell-EV treated cells. For Pre-miR-122 level, GAPDH was used as endogenous control (B) (mean+/− S.E.M). (C-D) Effect of Pre-miR-146a expressing macrophage derived-EV treatment on recipient hepatocytes. A schematic representation of the experiment has been described in panel C. Relative levels of miR-146a (D, left panel, p=0.0213, N=3), miR-122 (D, middle panel, p=0.0036, N=3) and Pre-miR-122 (D, right panel, p=0.0307, N=3) was measured by qRT-PCR in recipient Huh7 cells after pCI-neo and pmiR-146a transfected RAW264.7 cell derived-EV treatment for 24 h. U6 was used as endogenous control for miRNA normalization and miRNA levels in pCI-neo transfected cell-derived EV treated cells were used for normalization of miR-146a expressing cell-EV treated cells. For Pre-miR-122 level, GAPDH was used as endogenous control (D) (mean+/− S.E.M). (E) Effect of Pre-miR-146a overexpression on miR-122 level in hepatocytes. Relative levels of miR-146a (E, left panel, p=0.2251, N=3), miR-122 (E, middle panel, p=0.0003, N=4) and Pre-miR-122 (E, right panel, p=0.0279, N=4) were measured by qRT-PCR in Huh7 cells after pCI-neo & pmiR-146a expression in hepatocytes. U6 was used as endogenous control for miRNA normalization and miRNA levels of pCI-neo transfected cells was used for normalization of pmiR-146a transfected cells. For Pre-miR-122 level, GAPDH was used as endogenous control (mean+/− S.E.M). (F-H) Effect of EV treatment on Dicer1 level in naïve hepatocytes. Level of Dicer1 was measured by western blot in control and *Ld*-infected cell-EV treated Huh7 cells. β-Actin was used as loading control (F). Dicer1 level was also measured in Huh7 treated with EVs derived from pCI-neo and pmiR-146a transfected RAW264.7 cells. β-actin was used as loading control (G). Effect of Pre-miR-146a expression on Dicer1 in hepatocytes was also determined ( H) (I) Effect of Pre-miR-146a expression on signalling component proteins in hepatocytes. Changes in P-ERK1/2, P-P38 were determined by western blot analysis in pCI-neo and pmiR-146a transfected Huh7 cells. β-Actin was used as loading control (H, I). (J-K) Effect of NHA-Dicer1 overexpression on Dicer1 associated precursor miRNA level in cells expressing pre-miR-146a. Average Ct value of NHA-Dicer1 associated pre-miR-146a (J. left panel, p=0.0005, n=3) and pre-miR-122 level (J., right panel, p=0.2630, n=3, unpaired *t*-test) were determined by Real time PCR after HA immunoprecipitation from pCI-neo and pmiR-146a transfected Huh7 (J) (mean+/− S.E.M). NHA-Dicer1 was immunoprecipitated using anti-HA antibody and was detected in immunoprecipitated and input samples using anti-HA antibody (K). In all experimental data, ns: non-significant, *, ** and *** represent *P*-value of <0.05, <0.01 and <0.0001 respectively calculated by student paired *t-*test where not mentioned otherwise.

### Infected cell secreted EVs downgrade inflammatory response in neighbouring macrophage and induce macrophage polarization

While lowering of miR-122 levels in hepatocytes could be one of the mechanisms to reduce the proinflammatory response in the naïve macrophage, there must be additional mechanisms to restrict the production of inflammatory cytokines in the neighbouring naïve macrophage cells adjacent to the infected macrophages. To score the effect of infected cell-EVs on naïve macrophage cells, we treated RAW264.7 cells with EVs isolated from *Ld* infected RAW264.7 cells (Figure 6A). Interestingly, when the naïve RAW264.7 cells were pre-treated with *Ld* infected cell derived EVs, we documented a lesser responsiveness and relatively low proinflammatory cytokine production in the treated cells exposed to proinflammatory agents like LPS. Interestingly, the microvesicles isolated from the infected cells increase the production of cytokines in treated cells on LPS exposure (Figure 6B).

**Figure 6.**
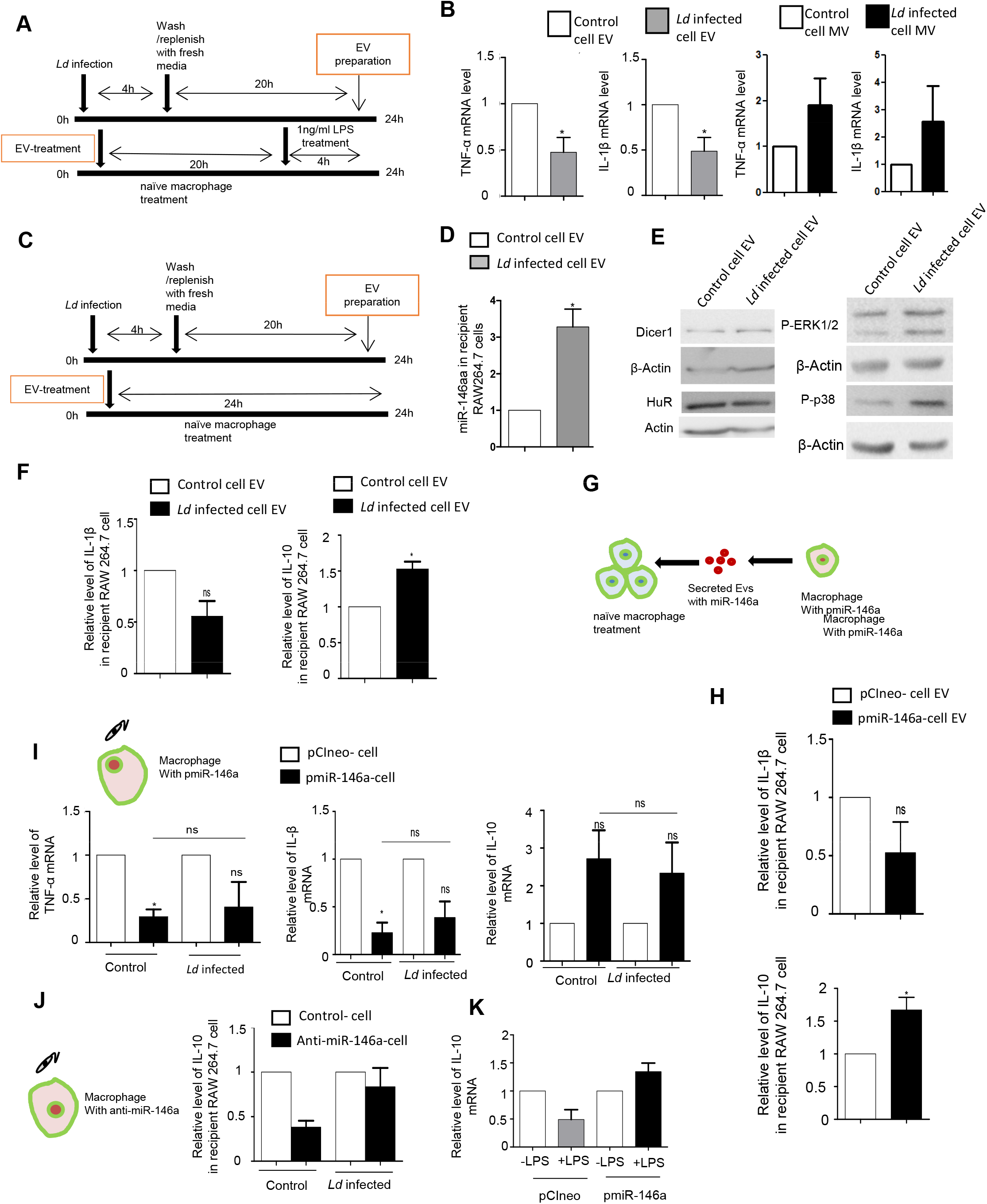
EV mediated transfer of miR-146a promotes ant-inflammatory response in recipient macrophages. (A-B) Effect of EV-treatment on LPS induced activation of recipient macrophages. A schematic representation of the experiment has been shown in the left (A). Relative cellular levels of TNF-α (B, left panel, p=0.0224, N=6), IL-1β (B, middle left panel, p=0.0280, N=5) mRNAs. TNF-α and IL-1β (B, right panels, N=2) were measured by qRT-PCR from Infected cell-EV treated or Micro Vesicles (MV)-treated macrophages after 4 h of LPS (1ng/ml) treatment.18s was used as endogenous control. Values in the non-infected cell-EV or MV treated set were used as units (mean+/− S.E.M). (C-F) Effect of infected cells derived EV-treatment on recipient macrophages. A schematic representation of the experiment has been shown (C). Relative level of miR-146a in recipient macrophage after 24h of EV treatment was measured by qRT-PCR and non-infected cell EV-treated cell miRNA level was set as unit. U6 was used as endogenous control (D, p=0.0184, N=4). Levels of Dicer1, HuR, P-ERK1/2, P-p38 were measured by western blot analysis in infected cell derive-EV treated RAW264.7 cells. β-Actin was used as loading control (E). Relative levels of IL-1β (F, left panel, p=0.0967, N=3) and IL-10 (F, right panel, p=0.0357, N=3) were measured by qRT-PCR in recipient RAW264.7 cells after 24h of EV treatment. GAPDH was used as endogenous control and values for non-infected cell EV treated cells were set as unit (F) (mean+/− S.E.M). (G-H) Effect of EV-mediated transfer of miR-146a on recipient macrophage. A schematic diagram has been given (G). Relative level of cytokine mRNA IL-1β and IL-10 were determined by qRT-PCR in recipient macrophage after treatment with EVs-derived from pCI-neo and pmiR-146a transfected macrophage (H, upper panel, p=0.2131,N=3 and lower panel, p=0.0406, N=4 respectively). GAPDH was used as endogenous control and values for control EV treated cells were set as units. (mean+/− S.E.M). (I) Effect of *Ld* infection on cytokine mRNA levels in miR-146a overexpressed macrophages. A schematic diagram of the experiment has been shown (I, upper panel). Relative cytokine mRNA level of TNF-α (I, left panel, p=0.0138, p=0.1740, p=0.7287, N=3), IL-1β (I, middle panel, p=0.0183, p=0.0684, p=0.4727, N=3) and IL-10 (I, right panel, p=0.1491, p=0.2417, p=0.7457, N=3) were measured by qRT-PCR from pCI-neo and pmiR- 146a transfected RAW264.7 cells after 24 h of *Ld* infection or no infection. GAPDH was used as endogenous control and values for pCI-neo control cells was set as a unit (I, unpaired *t-*tests were performed for comparison between pmiR-146a expressing non-infected control and infected set)(mean+/− S.E.M). (J) Effect of control and infected cell derived-EV treatment on recipient macrophages transfected with control or miR-146a inhibitor oligos. A schematic diagram has been given (J, left panel). Relative cytokine mRNA level of IL-10 was measured by qRT-PCR from miR-146a inhibitor transfected and EV treated RAW264.7 cells (J, right panel, N=2). GAPDH was used as endogenous control and values for negative control inhibitor transfected cells were set as units (mean+/− S.E.M). (K) Effect of LPS treatment on pmiR-146a expressing macrophages. Relative level of IL-10 was measured by qRT-PCR from control or pmiR-146a expressing cells with or without LPS exposure (1ng/ml). GAPDH was used as endogenous control and values for untreated control cells were set as units. (mean+/− S.E.M, N=2,). In all experimental data, ns: non-significant, *, ** and *** represent *P*-value of <0.05, <0.01 and <0.0001 respectively calculated by student paired *t-*test where not mentioned otherwise.

To score the effect of infected cell-EVs on polarization of naive macrophage, we treated the RAW264.7 macrophage with *Ld*-infected EVs (Figure 6C) and measured the amount of internalized miRNA (Figure 6D). With EV-treatment, we noted no change in Dicer1 level and a decrease in HuR expression with increased levels of p-P38 and ERK level (Figure 6E). These were accompanied by an increase in IL-10 expression and decrease in expression of proinflammatory cytokine IL-1β (Figure 6F). To conclude on casualty of EV-associated miR-146a with changed cytokine expression observed in *Ld*-infected cell derived EV-treated cells, we used EVs released by miR-146a expressing RAW264.7 cells and treated naïve RAW264.7 cells with miR-146a containing EVs. We noted an increase in IL-10 and a decrease in IL-1β after miR-146a containing EV-treatment also (Figure 6G and H). We expressed miR-146a in naïve and Ld-infected RAW264.7 cells and have noted a decrease in IL-1β and TNF-α expression and increase in IL-10 expression (Figure 6I). Interestingly, inhibition of miR-146a by anti-miR-146a reduced the expression of IL-10 both in naïve and Ld-infected RAW264.7 cells while the activation of cells with LPS have a reduced effect on anti-inflammatory IL-10 lowering, in miR-146a expressing cells. Both data, suggest a M2 polarization of miR-146a-treated macrophage that become refractory to immuno-stimulation by LPS (Figure 6I and J). To conclude on the contribution of *Ld*-derived factors associated with EVs-released by infected cells on naïve cell polarization and IL-10 production, we treated RAW264.7 cells with *Ld*-derived EVs and Lipophosphogylcan (LPG) isolated from the *Ld* membrane. We have noted no major change in the expression of major signalling component except an increase in p-P38 in *Ld*-derived EV treated cells (Supplementary Figure S7A, B and D). As reported earlier, we have also noted decreased TNF-α expression in both sets of treated cells while IL-1β only reduces in presence of Ld-derived EVs but not with *Ld* LPG [Supplementary Figure S7C and E,(de Carvalho et al, 2019)]. The IL-10 level decreased in both *Ld*-derived EVs and LPG treatment and therefore they cannot be the major player in observed increase in IL-10 in neighbouring macrophage cells (Supplementary Figure S7C and E). Taken together we conclude that the EV-associated miR-146a of Ld-infected cell EVs, have an anti-inflammatory effect on naïve macrophage cells that is associated with an increase in IL-10 production in treated cells while the LPS responsiveness as well as proinflammatory cytokine level remain reduced in miR-146a enriched RAW264.7 cells.

### HuR is required for miR-146a mediated IL-10 induction

From the data discussed so far it is becoming more likely that miR-146a control IL-10 level in macrophage cells. How does miR-146a ensure a high IL-10 expression in treated macrophage to polarize them to have an anti-inflammatory response pathway- a prelude to the infection niche establishment? HuR is an important post-transcriptional gene regulator in mammalian macrophage that is essential for activation and expression of pro-inflammatory cytokines in macrophage exposed to LPS. Activated macrophage also showed increased expression of HuR with LPS-treatment time (Goswami et al, 2020). Interestingly, expression of HA-HuR also enhances the expression of pro-inflammatory cytokines in untreated macrophages and could get them in activated state (Fig 7A and B, Goswami et al 2020). Therefore, in order to have the strong anti-inflammatory effect of miR-146a, the HuR-mediated pro-inflammatory effect should be neutralized or countered by the transferred miR-146a from infected cells. *Leishmania* could downregulate HuR in mammalian macrophage cells (Supplementary Figure 5H). However, it is possible that the extracellular miR-146a released by the infected cells could bring down the effect of HuR in non-infected cells present in the infection niche to ensure an overall anti-inflammatory pathway prevalent in all immune cells present there. miR-146a mediated downregulation of HuR has been reported previously (Cheng et al, 2013). Does miR-146a counter the action of the HuR to stop inflammatory responses in macrophage cells? In miR-146a containing EV-treated cells, we have noted that the expression of HuR could not enhance the IL-1β or TNF-α expression in RAW264.7 cells. Surprisingly, an increase in IL-10 expression in HuR expressing cells was detected when treated with miR-146a containing EVs (Figure 7C-D). Inversely siHuR treatment causes reduction in IL-10 expression in cells treated with miR-146a containing EVs (Figure 7E-F). Testament of miR-146a mimic decreases HuR levels as well as HuR induced increase in TNF-α and IL-1β expression while IL-10 level increases in presence of HA-HuR when miR-146a is expressed in RAW264.7 cells (Figure 7G-K). How does HuR ensure the high IL-10 level? It is known that miR-21 negatively regulates IL-10 (Wang et al, 2017) and HuR is also known to interact and sponges out miR-21 to inactivate it in mammalian cells (Poria et al, 2016).. We have noted redu.ction in miR-21 level in HuR expressing cells and siHuR treatment prevent miR-21 lowering. We have also noted increased miR-21 export in HuR expressing cells that can confirm that HuR by exporting miR-21 helps the IL-10 expression in cells treated with miR-146a (Figure 7L-M). Thus, HuR by upregulating IL-10 level also ensure the strong anti-inflammatory polarization in miR-146a containing EV-treated cells. Interestingly HuR is getting decreased in miR-146a expressing cells to balance the HuR mediated stabilization of cytokine mRNAs like TNF-α and IL-1β to further stimulate the anti-inflammatory response. Therefore it is the balanced expression of HuR and miR-146a that determine the polarization status of the macrophage. In miR-146a negative situation, expression of HuR promotes M1 polarization (Goswami et al, 2020), while with increasing concentration of miR-146a the HuR mediated miR-21 export contribute in IL-10 expression. High IL-10 down regulates HuR level to ensure the checked expression of pro-inflammatory cytokines and its own expression in a feedback effect (Rajasingh et al, 2006).

**Figure 7.**
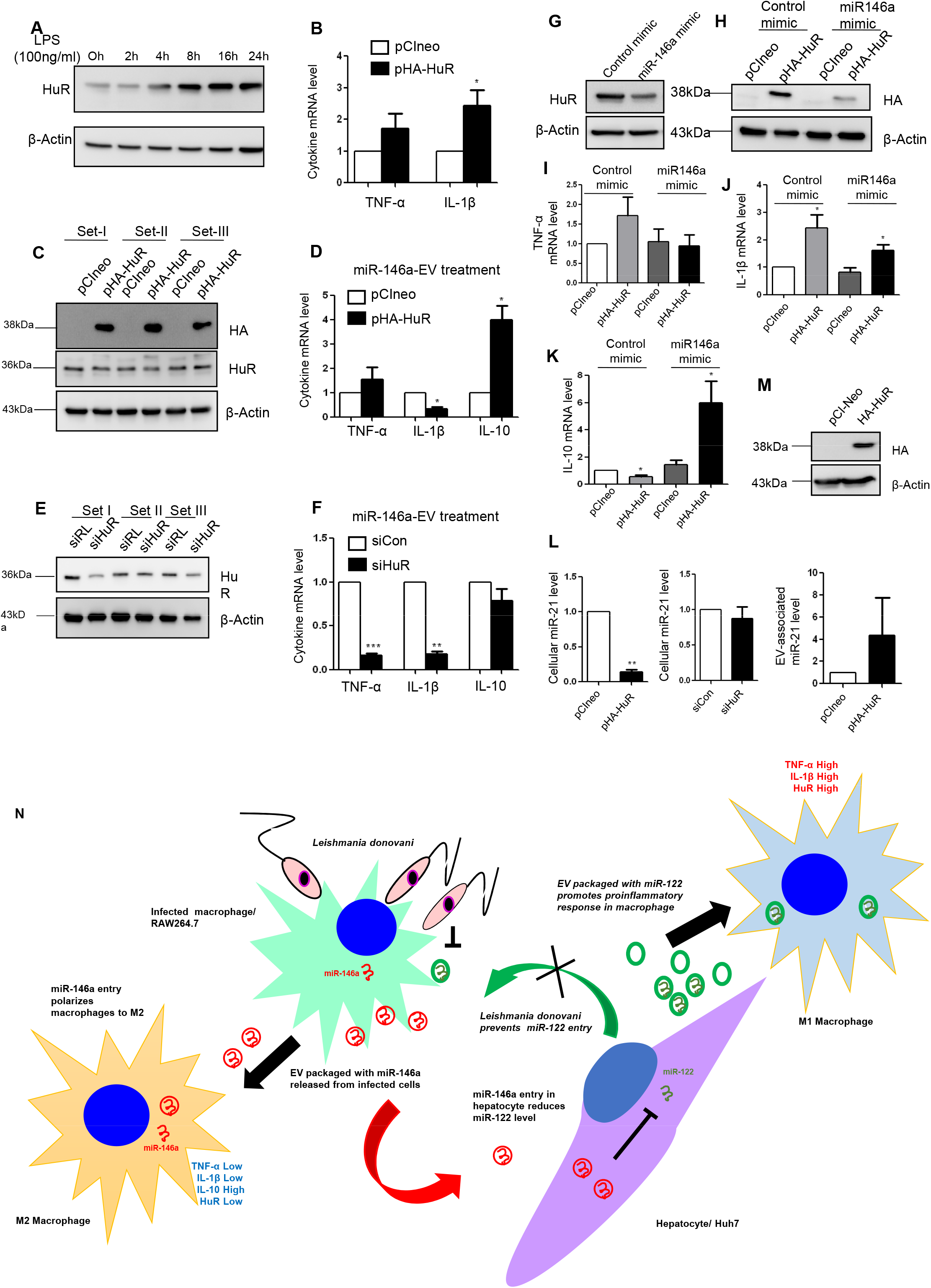
Cross-talk between HuR and miR-146a determines the fate of cytokine response in macrophage cells. (A) LPS treatment for different time points in RAW264.7 cells. The cell lysates were immunoblotted for HuR. β-Actin was used as loading control. (B) qRT-PCR was done for pro-inflammatory cytokine mRNAs (TNF-α and IL-1β) in pCI-neo and pHA-HuR transfected cells (mean +/− sem; n=3), (C, D) EVs isolated from pmiR-146a transfected RAW 264.7 cells were treated to recipient RAW264.7 cells expressing pCI-neo or pHA-HuR. Recipient cells treated with pmiR-146a expressing cell-EVs were immunoblotted for HA and HuR, β-Actin was used as loading control (C). Real time PCR was done for cytokine mRNAs (TNF-α, IL-1β and IL-10) in pCI-neo and pHA-HuR transfected recipient cells treated with pmi-146a-EVs, GAPDH was used for normalization (mean +/− sem; n= 3; p-values: 0.3952, 0.0134, 0.0352 respectively) (D). (E, F) EVs isolated from pmiR-146a transfected RAW 264.7 cells and treated to recipient RAW 264.7 cells depleted of HuR. Recipient cells treated with pmiR-146a-EVs were immunoblotted for HuR to check for proper silencing and β-Actin was used as loading control (E). Real time PCR was done for cytokine mRNAs (TNF-α, IL-1β and IL-10) in siCon and siHuR containing recipient cells treated with pmiR-146a-EVs, GAPDH was used for normalization (mean +/− sem; n= 3; p-values: 0.0004, 0.0015, 0.2641respectively(F). (G-K) Effect of HA-HuR expression in RAW264.7 cells transfected with pre-miR control mimic or pre-miR146a mimic. Transfection was done either with pCIneo (control plasmid) or pHA-HuR. Immunoblotting the cell lysates for HuR using β-Actin as loading control (G). Immunoblotting the respective cell lysates for HA to check for proper overexpression of HuR, β-Actin used as loading control (H). Real time PCR was done for TNF-α, GAPDH served as control. Values obtained for pCIneo and pre-miR control mimic co-transfected sets were taken as units (mean +/− sem; n= 6; p-values: 0.2048, 0.8587, 0.8620 respectively). (I) Real time PCR was done for IL-1β, GAPDH served as control. Values obtained for pCIneo and pre-miR control mimic co-transfected sets were taken as units (mean +/− sem; n= 6; p-values: 0.0467, 0.3453, 0.0375 respectively) (J). Real time PCR was done for IL-10, GAPDH served as control. Values obtained for pCIneo and pre-miR control mimic co-transfected sets were taken as units (mean +/− sem; n= 6; p-values: 0.0239, 0.2710, 0.0267 respectively) (K). (L, M) Relative level of miR-21 in HuR overexpressed RAW264.7 cells. Real time PCR for miR-21 level in RAW264.7 cells expressing HA-HuR and transfected with pre-miR146a mimic. U6 snRNA served as control (mean +/− sem; n= 3; p-value: 0.0015) (L, left panel). Real time PCR for miR-21 level in recipient RAW264.7 cells depleted of HuR which were treated with miR-146a containing-EVs. U6 snRNA served as control (mean +/− sem; n= 3; p-value: 0.5259) (L, middle panel). Real time PCR of EV-associated miR-21 level from EVs of RAW264.7 cells expressing HA-HuR. EV marker protein Flotillin-1 was used for normalization of miR-21 level (mean +/− sem; n= 3; p-value: 0.4313) (L, right panel). Immunoblotting the cell lysates with HA using β-Actin as loading control in pCIneo or pHA- HuR transfected RAW264.7 cells (M). (N) The model depicts the protective role of extracellular vesicle associated miRNAs in *Ld* infection. The left half shows that *Ld* infection triggers EV mediated secretion of miR-146a from macrophage which when transfer to naïve macrophage prevent parasite internalization whereas the right half represent that when miR-146a get delivered to naïve hepatocyte via EV it reduces miR-122 level which in turn prevent *Ld* infection in macrophage due to its pro-inflammatory role. For all the experimental data; * signifies p-value <0.05, ** signifies p-value <0.01 and *** signifies p-value <0.0001. Statistical analysis was done using paired student’s t tests for all the experiments, unless mentioned otherwise. Positions of molecular weight markers are represented in the immunoblots used in the different panels.

### miR-146a acts as a balancer of IL-10 production and *Ld* infection

Does miR-146a uptake via EV affect the Ld-infection process? We have measured the number of parasites internalized in the treated macrophage after *Ld*-infected cell-EVs or miR-146a containing-EVs treatment. We have performed quantification and noted reduced entry of *Ld* after the miR-146a positive or infected cell EV-treatment (Supplementary Figure 8A-D). Lower expression level of *Leishmania* amastigote specific gene amastin also confirmed reduced levels of *Ld* entry after the miR-146a containing-EV or infected cell-EV treatment of RAW264.7 cells (Supplementary Figures S8E-G). miR-146a expression in RAW264.7 cells also reduces the *Ld* infection process. We have used single cell infection level examination to conclude on preferential reduction of *Ld* internalization in cells expressing miR-146a (Supplementary Figures S8H). Infection level may be downregulated due to a problem in signalling pathways or downstream factors. We have noted an increased p-p38 level in miR-146a expressing macrophage that signifies the existence of a possible counteractive pathway that usually gets reduced in cells infected with *Ld* where a p38/MAPK downregulation has been noted (Supplementary Figures S8M). The decreased *Ld* infection could have also been attributed by a possible reduction in the endocytosis process itself. However, in a latex bead internalization assay, we did not document any reduction in latex bead internalization after miR-146a containing EV or infected cell EV treatment of RAW264.7 cells. Expression of miR-146a also doesn’t affect the latex bead entry process in RAW264.7 cells (Supplementary Figure 8I-L).

## Discussion

Extracellular vesicles are known for their role in intercellular communication. They play an important role in immune response during any pathological condition and also help to establish tumor microenvironment (Thery et al, 2002). In disease condition, EVs either can help in progression of the disease or in providing protection against the disease. There are many studies where it has been reported that exosomes from *Mycobacterium* infected cells can trigger proinflammatory response in noninfected cells (Bhatnagar et al, 2007). *Leishmania* parasites release exosomes as a vehicle to transport proteins to the host cell for immunosuppressive action (Silverman et al, 2010). *Leishmania* protein GP63 was found to be transported to neighbouring hepatocytes via secreted exosomes that target Dicer1, a processor of pre-miRNAs to downregulate expression of liver specific miR122 (Ghosh et al, 2013). *Leishmania* parasite secretes exosomes within sandfly midgut and these exosomes are part of the sandfly inocula during bite which help in pathogenesis via overproduction of IL-17a in target cells (Atayde et al, 2015). Gioseffi et al. in recent time have reported *Leishmania* infected EVs as carrier of large number of Leishmanial and host proteins (Gioseffi et al, 2020). Among the proteins exported out, they have identified that a *Leishmania* homolog of mammalian angiogenesis promoting factor, vasohibin is found to induce endothelial cells to release angiogenesis promoting factors and thereby helping in promotion of lesion vascularization during infection.

miRNAs, being the regulator of expression of several cytokines, are important players in determining the fate of immune cells and in particular have a major role in buffering the immune activation processes by regulating the expression of several cytokines directly or indirectly (Lindsay, 2008; Testa et al, 2017). miR-146a and miR-155 are the two most important players that are explored for their potential role as immune checkpoint regulators in the mammalian system. While miR-155 is a proinflammatory miRNA, miR-146a acts oppositely to balance the immune activation process (Testa et al, 2017). In the infection context, the pathogens need to target the miRNAs pathways to control the inflammation level. The *Ld* parasite has shown previously to control expression of miRNAs in a negative manner in infected cells. By downregulating HRS and depolarizing mitochondria in the host, *Ld* affects the miRNA recycling process to enhance the miRNA content of the infected cells. However, the accumulated miRNAs are largely non-functional as they found to be accumulated with endosomal fraction and they fail to recycle for new round of repression on ER attached polysomes (Bose et al, 2020). The miR-146a that also increases with *Ld* infection gets exported to neighbouring cells. The miR-122 is another miRNA in hepatocytes that *Ld* needs to control to ensure lower cholesterol biogenesis in the liver cells essential for parasite survival. *Ld* by targeting Dicer1 in hepatocytes control the expression of miR-122 in infected liver tissue (Chakrabarty & Bhattacharyya, 2017; Ghosh et al, 2013). Our results show that miR-146a, transferred from infected macrophage cells, could also have a negative role to play on miR-122 expression in mouse liver. Reciprocally miR-122 can get transferred from activated or stressed liver cells to resident macrophage to get them activated (Ghosh et al, 2015; Mukherjee et al, 2016). The parasite uses a secondary mechanism of lowering the miR-122 in liver cells to ensure reduced transmission of the proinflammatory miR-122 to macrophages to prevent the expression of inflammatory cytokines there. EV-mediated crosstalk between the macrophages and hepatocytes fosters an environment of cell-to-cell communication to take place. However, in the liver, it is known that the EVs released from LPS treated THP-1 macrophages stimulated the proliferation of hepatic stellate cells, while miR-103-3p present in the EVs isolated from macrophages could get transferred to the hepatic stellate cells and promotes its growth (Chen et al, 2020).

MicroRNAs are the small noncoding RNAs that are involved in post-transcriptional & translational regulation of various genes. Thus, altered expression of these miRNAs are one of the main reasons for various pathogenic conditions (Bartel, 2018). Earlier reports suggested upregulation of miR-146a/b during Salmonella infection (Ordas et al, 2013) Upregulation of miR-155, miR-146a/b and downregulation of miR-20a, miR-191, miR-378 in *Mycobacterium avium* infected condition were also reported (Curtale et al, 2019; O’Connell et al, 2012). miR-155 favors the pro-inflammatory environment while miR-146a favours the anti-inflammatory environment (Lindsay, 2008). Recently miR-146a mediated M2 polarization during *Leishmania donovani (Ld)* infection has been reported. The investigation revealed that *Leishmania donovani (Ld)* infection upregulates miR-146a expression which favoured parasite survivability in hosts by maintaining the anti-inflammatory environment. Inhibition of miR-146a lowered parasite burden in infected Balb/C mice as well as anti-inflammatory environment (M2 phenotype). Interestingly this upregulation of miR-146a during *Ld* infection is regulated by Super Enhancer components like BET bromodomain 4 (BRD4) and P300 (Das et al, 2021).

The polarization of macrophage is contributed by several factors including specific activation of signalling component that favours either the TLR4-p38/MAPK-NF-kB mediated activation of pro-inflammatory cytokine expression or it may be an inhibitory circuit involving ERK dependent inactivation of the pro-inflammatory cytokines with concomitant increase in expression of anti-inflammatory cytokine like IL-10 (Mukherjee et al, 2021). The parasite seems to use the infected cell-EVs packed with miR-146a to transmit the anti-inflammatory signals to the naive neighbouring macrophage to polarize them to the M2 stage. It is not known whether the polarization is reversible, but subsequent treatment of miR-146a treated cells with LPS suggest the refractory nature of the recipient macrophage to inflammatory response. Thus miR-146a containing EVs could be considered as a good immune modulator and use of that could be important to control the inflammation in different physiological contexts like in prevention of viral infection related cytokine storm (Arisan et al, 2020) or inflammation associated with tumour (Rupaimoole et al, 2016). The cross communication of miR-146a containing EVs to other non-infected tissue is another important aspect. Do EVs with miR-146a get transferred via the bloodstream to the spleen and do they have any effect on subsequent establishment of infection and T-cell polarization observed in the spleen of infected animals? These are important questions for future exploration.

HuR is a RNA binding protein with immense functional diversity. HuR is known to stabilize the mRNAs with AU-rich elements in their 3’ UTR and thus have a role in the inflammation process as most of the pro-inflammatory cytokines like TNF-α or IL-1β bear the AU-rich elements in their 3’ UTR (Srikantan & Gorospe, 2012). In this report, we have detected a non-canonical role of HuR in stabilization of IL-10 mRNA. The anti-inflammatory IL-10 mRNA is regulated by miR-21 negatively (De Melo et al, 2021), the suppression of miR-21 activity by HuR has been reported previously (Poria et al, 2016). In the context of miR-146a induced expression of IL-10, HuR acts synergistically to remove miR-21 from the cell to boost the anti-inflammatory IL-10 production in macrophages. Interestingly, in the context of an anti-inflammatory environment the miRNA depressor role of HuR predominate to ensure expression of miRNA-repressed cytokine mRNAs while in the pro-inflammatory context the mRNAs stabilizer role of HuR is important to ensure proinflammatory cytokine expression there.

## Materials and Methods

### Cell culture, peritoneal macrophage isolation and parasite infections

Human hepatic Huh7, mouse hepatic Hepa1-6 and Mfn2 wildtype and knockout mouse embryonic fibroblasts cells (WT/Mfn2^−/−^) were cultured in Dulbecco’s Modified Eagle’s Medium (Gibco-BRL) and 10% FBS (heat inactivated Foetal Bovine Serum). For EV free growth media, 10% EV depleted FBS (made by ultracentrifugation of FBS at 1,00,000X g for 4 hours) was added to DMEM.

In case of RAW 264.7 macrophage cells, RPMI 1640 medium (Gibco) with 2mM L-glutamine and 0.5% β-Mercaptoethanol along with 10% FBS was used. Primary murine peritoneal macrophages were isolated from BALB/c mice subjected to intraperitoneal injection of 1.5ml of 4% starch solution. Cold 1X PBS was used to lavage or wash the peritoneal cavity in order to isolate the peritoneal macrophages the following day which were then pelleted and plated. These cells were also cultured in RPMI 1640 like the RAW macrophages. EV isolation of Peritoneal Epithelial Cells (PEC) was done from a 60 mm plate. All experiments with PEC were done after 48 hours of isolation (Goswami et al, 2020). For macrophage activation, LPS dose of 1ng/ml was used for 4 hours. RAW264.7 and Huh-7 cells were stimulated with LPS from *Escherichia coli* O111:B4 (Calbiochem).

### Animal experiments

BALB/c male mice (4-6 weeks) were divided into four groups (three animals each) of either uninfected or infected and treated or untreated with EVs. Two groups (six animals) were infected with 1 × 10^7^ promastigotes by intracardiac puncture and maintained for 30 days. On the 30th day, EVs were injected into six animals (three infected and three uninfected with parasites) while 1X PBS was injected in the remaining six animals (three infected and three uninfected). For EV isolation, Hepa1-6 cells were transfected with miR-122 expressing plasmid and approximately 5 x 10^9^ EVs (measured by Nanoparticle Tracker Nanosight NS300) isolated from these transfected cells were suspended in 1X PBS and injected into the tail vein. The animals were sacrificed on the 31^st^ day for Kupffer cell isolation.

All the experiments were done in accordance with the National Regulatory Guidelines issued by the Committee for the Purpose of Supervision of Experiments on Animals, Ministry of Environment and Forest, Govt. of India. In agreement with the Institutional Animal Ethics Committee the animal experiments were done. The BALB/c mice were kept under controlled conditions (temperature 23 ± 2°C, 12 hour/12-hour light/dark cycle) in individually ventilated cages.

Kupffer cell isolation was done from all the mice as mentioned previously (Aparicio-Vergara, 2017) with minor modifications. The animals were anaesthetized and livers were perfused with HBSS via the hepatic portal vein with an incision in the inferior vena cava until they turned bloodless and then digested with a Liver Digest Medium. Livers were then excised, minced and filtered through a 70μm cell strainer. The resultant single cell suspension was centrifuged at 50X g for 5 minutes to pellet and store the hepatocytes. The supernatant was then collected and loaded onto a Percoll gradient of 25% and 50%. It was centrifuged at 1200X g for 30 minutes at 4°C without brakes. The interphase containing the Kupffer cells was collected and washed twice with PBS and the pellet was stored. The Kupffer cell pellet and hepatocyte pellet were subjected to RNA isolation by TriZol reagent.

For detection of purity of isolation, the Kupffer cells of all the four groups were compared with the respective hepatocytes by quantification of mRNA levels of C-type Lectin Domain Family 4, Member F (Clec4f), a Kupffer Cell marker and hepatocyte marker, Albumin. The kinetoplast DNA levels were also quantified in the four groups to detect the level of leishmanial infection.

### Parasite culture and infection to macrophage cell line

*Leishmania donovani (Ld)* strain AG83 (MAOM/IN/83/AG83) was obtained from a Indian Kala-azar patient and was maintained in golden hamsters (Chakrabarty & Bhattacharyya, 2017). Amastigotes were obtained from infected hamster spleen and transformed into promastigotes. Promastigotes were maintained in M199 medium (Gibco) supplemented with 10% FBS (gibco) and 1% penstrep (Gibco) in 22°C. RAW 264.7 cells or PEC were infected with stationary phase *Ld* promastigotes of 2^nd^-4^th^ passage at a ratio of 1:10 (cell: *Ld*) for 6h, 16h or 24 h depending upon the experiment. (Goswami et al, 2020).

### Plasmid constructs & transfection

pCIneo Precursor miR-146a (pmiR-146a) and HA-HuR plasmids were transfected using Fugene HD (Promega) for RAW264.7 cells as described previously (Goswami et.al, 2020). For Huh7 cells, all transfections of plasmids were performed using Lipofectamine 2000 reagent (Life technologies) according to the manufacturer’s protocol. For miRNA inhibitor (anti-miR-146a) experiments, 30 picomoles per well was transfected using RNAimax reagent in RAW264.7 cells according to manufacturer’s protocol. Transfection of Negative control inhibitor was used for controls. For co-transfection of plasmid and pre-miR mimic, Lipofectamine 2000 (Invitrogen) was used. 50pmoles of pre-miR mimic were transfected per well of a 12-well plate. siRNA transfection was performed using RNAi max (Invitrogen) following manufacturer’s instructions. RAW 264.7 cells were transfected with 50 pmoles of siRNA per well of a 12-well plate. EV treatment was given to the recipient cells 48h after siRNA transfection and the cells were harvested 24h after EV treatment. Donor RAW 264.7 cells were transfected with 2 μg pmiR-146a plasmid per well of a 6-well plate using Fugene HD.

### Extracellular vesicle (EV) isolation & characterization

RAW264.7 cells were plated in a 90 mm plate and given infection with stationary phase parasite at about 80% confluency and kept for 6h. After 6h of infection media was discarded and cells were washed with PBS to remove extracellular unattached parasites and cells were replenished with fresh RPMI1640 medium supplemented with 10% EV depleted FBS and 1% penstrep for 18h. Total infection time was 24h for EV isolation. After 24h the supernatant was collected for EV isolation. Briefly, the supernatant first centrifuged at 3,000xg for 15 minutes at 4^0^C followed by 30 minutes centrifugation at 10,000xg at 4^0^C. For all EV isolation experiments, supernatant was passed through a 0.22μm filter unit followed by ultracentrifugation on a 30% sucrose cushion at 1,00,000xg for 70 minutes. The supernatant was discarded leaving a medium layer of interface containing EVs behind, which was then washed with 1X PBS at 1,00,000xg for 70 minutes again. The pellet then was resuspended in PBS for NTA or 1X Passive Lysis Buffer (PLB) for RNA or protein estimate (Promega). For EV associated RNA and protein, the pellet was resuspended in PLB. One-third was used for total RNA isolation using TRIzol LS reagent (Invitrogen) following the manufacturer’s protocol. Two-third of the PLB sample was used for protein precipitation using methanol precipitation. In brief, for 200 µl of PLB sample 640 µl of sterile water, 480 µl of methanol (Merck) and 160 µl of chloroform (Sigma) were added and vortex well followed by centrifugation at 20,000xg for 5 min at room temperature. Then the supernatant was discarded and 300ul of methanol was added. The mixture was vortexed well and again centrifuged for 30 mins at 20,000xg 4°C. The supernatant was discarded. The pellet was air dried for 10 mins and then dissolved in 1x SDS Dye and heated at 95°C.

For experiments with EV treatment, culture supernatant first centrifuged at 3,000xg for 15 minutes at 4^0^C followed by 30 minutes centrifugation at 10,000xg and filtered pass through 0.22 µm following ultracentrifugation at 1,00,000xg for 70 minutes. The pellet was then resuspended in RPMI-1640 medium supplemented with 10% EV depleted FBS and 1% penstrep for further use.

For Nano-particle tracking analysis (NTA) the control and infected RAW264.7 cell derived EV pellet was resuspended in 300ul of sterile PBS and resuspended well. Then diluted 10 fold in PBS and 1 ml of diluted sample was used for NTA (Nanosight NS300).

### EV treatment of recipient cells

After EV isolation the pellet was resuspended in fresh RPMI1640 supplemented with 10% EV depletede FBS and 1% penstrep followed by filtration through 0.22 µm filter unit. The EVs were then added to the recipient macrophage or hepatic cell line for 24h. Post LPS treatment was given at a dose of 1ng/ml for 4h after 20h of EV-treatment to macrophages. For treatment EVs isolated from 8×10^6^ cells (one 90mm plate) were added to 2.4×10^5^ recipient cells (per well of a 12 well plate) approximately. For estimation of parasite infection after EV treatment, parasites were added to EV-treated cells for 6 h.

### Optiprep^TM^ Density gradient ultracentrifugation

For cell fractionation using 3%-30% iodixanol gradient (optiprep gradient) 1.6×10^7^ cells were used as described earlier (Mukherjee et al, 2016). Optiprep^TM^ (Sigma-Aldrich) solution was used to prepare 3%-30% gradient. In brief, the cell pellet was homogenized in a buffer with glass Dounce homogenizer. The lysate was centrifuged at 1,000xg twice and the supernatant was loaded on the gradient and centrifuged at 36,000 rpm in a SW60Ti rotor for 5h. Approximately 400 µl of 10 fractions were collected for RNA and protein.

### Total RNA isolation & quantification of miRNA and mRNA

Total RNA extract from experimental samples using TriZol reagent (Invitrogen) for cell and TriZol LS (Invitrogen) for fraction according to manufacturer’s protocol. All miRNA and mRNA quantification were done as described previously (Basu & Bhattacharyya, 2014; Mukherjee et al, 2016). In brief, miRNA estimation was performed with 100ng of RNA for cellular and EV samples using specific primers for human miR-146a, mouse miR-155, human miR-122, human miR-21, human miR-16, miR-125b and U6 snRNA was used as endogenous control. For the assay, one-third of the reverse transcription mix was subjected to PCR amplification with TaqMan® Universal Master Mix No AmpErase (Applied Biosystem) and respective TaqMan® reagents for target miRNAs. Samples were analyzed in triplicates for each biological replication. The comparative C_t_ method which involved normalization by U6 snRNA was used for quantification (Mukherjee et al, 2016).

For mRNA, 200 ng of total RNA was used for estimation based on SYBR Green based real-time cDNA estimation using specific primer for target genes. The comparative C_t_ method which involved normalization by GAPDH or 18s rRNA was used for quantification of mRNA (Goswami et al, 2020). The details of Primers, miRNA assays are provided as part of Supplementary Table S4-5.

### Western blot

Western blot analyses were performed as described elsewhere (Mazumder et al, 2013). Imaging of the blots was performed using an UVP BioImager 600 system equipped with VisionWorksLife Science software (UVP) V6.80 which was also used for quantification. Details of all antibodies used for Western blot and immunofluorescence experiments are summarized in Supplementary Table S6

### Immunofluorescence

For internalization, *Ld* parasites were stained with 1μM carboxyfluorescein succinimidyl ester (CFSE) dye (green) for 30 mins at 22^0^C followed by PBS wash thrice and then resuspended in medium before adding to the cell. Cells were fixed with 4% paraformaldehyde (PFA) for 20 minutes after three PBS wash. Nuclei were stained with DAPI. For calculating the percentage of infected cells, a minimum number of 100 cells were counted. Cells were observed under the Zeiss LSM800 confocal microscope.

### Polysome Isolation

For polysome isolation approximately 8×10^6^ cells were used as starting materials. After 24h of infection, cells were scraped in PBS and pellet down at 600xg for 5 min at 4 °C. Cell pellet was collected and incubated in 1X lysis buffer (10mM HEPES, 25mM KCL, 5mM MgCl2, 1mM DTT, 5mM VRC, 1X PMSF, Cycloheximide 100 µg/ml, 1% Triton X-100, 1% sodium deoxycholate) for 15 min at 4°C followed by centrifugation at 3,000xg for 10min at 4°C. Then the supernatant was collected and centrifuged at 20,000xg for 10 min at 4^0^C. The supernatant was collected and loaded on 30% sucrose cushion in gradient buffer and centrifuged at 31200 rpm for 1 hour in 4 °C in a SW61Ti rotor. After centrifugation the non-polysome fraction were collected and mixed rest of the solution followed by centrifugation at 31,200 rpm for 30 min at 4 °C in a SW61Ti rotor. The pelleted polysome was suspended in polysome buffer (10mM HEPES,25mM KCL, 5mM MgCl2, 1mM DTT, 5mM VRC, 1X PMSF) and kept for RNA isolation and western blot (Ghoshal et al, 2021).

### LPG extraction and treatment

LPG isolation was performed as mentioned earlier (Goswami et al, 2020). In brief, 2×10^8^ stationary phase parasites (AG83) were used for LPG extraction. The pellet was resuspended in 2 ml of Chloroform: Methanol: Water mixture (1:2:0.5 v/v) followed by vortex and sonication for 10 seconds thrice at 30 seconds interval and incubate in mutarotator at room temperature for 2h. Then the lysate was centrifuged at 4,000xg for 30 mins at 4°C. The pellet was used for extraction in 9% 1-Butanol in water. After vortex and sonication as before, pellet was incubated for 1h at room temperature in mutarotator. Second extraction was done with the pellet in 9% 1-Butanol in water again after centrifugation at 4,000xg for 30 mins at 4°C. The supernatants of both extractions were pooled for lyophilization. The lyophilized LPG was resuspended in 1X sterile PBS before cell culture treatment. For treatment, 1:50 (cell: LPG) dose was used for different time points.

### Mass spectrometry

For proteomics sample preparation, exosomes from two 90 mm were pooled and protein was precipitated with 100% chilled acetone in -20°C overnight. The protein precipitated was then washed with 80% chilled acetone followed by another wash with 40% chilled acetone. The pellet then air dried for 5-10 mins to remove excess acetone at room temperature and resuspended in 50mM ammonium bicarbonate (AmBic) followed by sonication thrice. 85mM DTT was then added and the sample was heated at 60 °C for 1h. Sample was then incubated in dark with 55mM Iodoacetamide (IAA) at RT for 30 mins. Trypsin(1µg/µl) digestion was done to the sample for overnight at 37 °C. the reaction was stopped with 100% formic acid (5% of total volume) and snap freeze until lyophilization. Lyophilized sample was used for proteomics and run in Orbitrap analyzer.

### Latex bead phagocytosis experiment

Red fluorescent labelled Latex bead (Fluorescent Red, Sigma-Aldrich) was diluted in RPMI-1640 medium and added to macrophages at a ratio of 1:10 (cell:bead) for 6hour. Cells were fixed with 4% PFA as described earlier and mount in DAPI. Cell cytoskeleton was stained with Phalloidin-488. Red latex bead phagocytosis was imaged in ZeissLSM800 confocal microscope.

### Statistical analysis

All graphs and statistical significance were calculated using GraphPad Prism 5.00 (GraphPad, San Diego). Experiments were done for minimum three times to get *P*-value, unless otherwise mentioned. Paired and unpaired Student’s *t*-tests were used for determination of *P*-values. The sample size was chosen by convenience and no exclusion criteria was used.

### Ethics statement

Balb/C mice and Syrian golden hamsters were obtained from CSIR-Indian Institute of Chemical Biology animal house facility. All experiments were performed according to the national regulatory guidelines stated by the Committee for the Purpose of Supervision of Experiments on Animals, Ministry of Environment and Forest, Govt. of India. All animal experiments were performed with prior approval of the Institutional Animal Ethics Committee.

## Acknowledgement

We acknowledge Witold Filipowicz for different plasmids constructs. SNB and KM acknowledge CSIR-Indian Institute of Chemical Biology (IICB) for infrastructural support.

## Funding

SNB is supported by The Swarnajayanti Fellowship and and a High Risk High Reward Grant from Dept. of Science and Technology, Govt. of India, and KM and SNB both received support from CSIR. SG received support from University Grant Commission, India. BG, IB, SC, SB, AG received their support from CSIR, India. The funders had no role in study design, data collection and analysis, decision to publish, or preparation of the manuscript.

## Author contributions

SNB and KM conceptualize and designed the study, interpreted the data and prepared the draft manuscript. SG, BG, and IB did majority of the experiments, did analysis and prepared the final data. They were supported by AG, SC and SB. All authors read and approved the final version of the manuscript.

## Supplementary Information

### Supplementary Figure Legends

**Figure S1.**
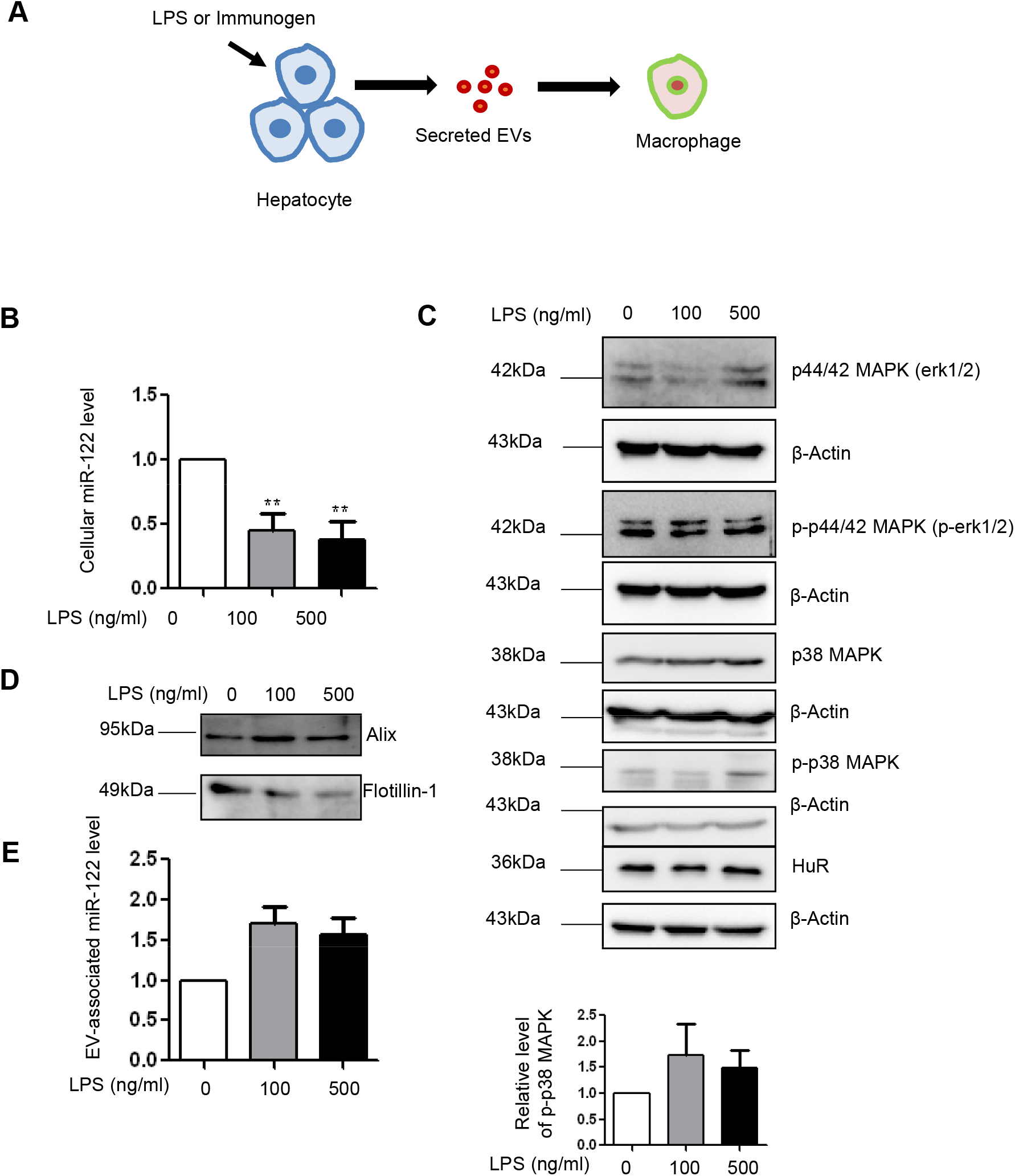
LPS stimulated hepatocytes exports out miR-122 through EVs. (A) Schematic representation of LPS stimulation of Huh7 cells. LPS stimulation of 100ng/ml and 500ng/ml were given to Huh7 cells for 24hrs. (B) Changes in cellular miR-122 levels in LPS stimulated Huh7 cells, U6 snRNA used as control (mean +/− sem; n= 6; p-values: 0.0092, 0.0070 respectively). (C) Immunoblots for ERK, P-ERK, p38, P-p38 and HuR were checked in cellular lysates of LPS stimulated Huh7 cells. β-Actin was used as loading control (upper panel). Quantification of relative level of P-p38 normalized with β-Actin (lower panel). (D) Immunoblots for confirming the presence of EV marker proteins Alix and Flotillin1 in EVs isolated from control and LPS stimulated Huh7 cells. (E) Quantification of miR-122 levels in EVs isolated from LPS stimulated Huh7 cells. EV protein markers were used for normalization of miR-122 levels (mean +/− sem; n= 5; p-values: 0.0809, 0.1085 respectively). For all the experimental data; * signifies p-value <0.05, ** signifies p-value <0.01 and *** signifies p- value <0.0001. Statistical analysis was done using paired student’s t test for all the experiments, unless mentioned otherwise. Positions of molecular weight markers are represented in the immunoblots used in the different panels.

**Figure S2.**
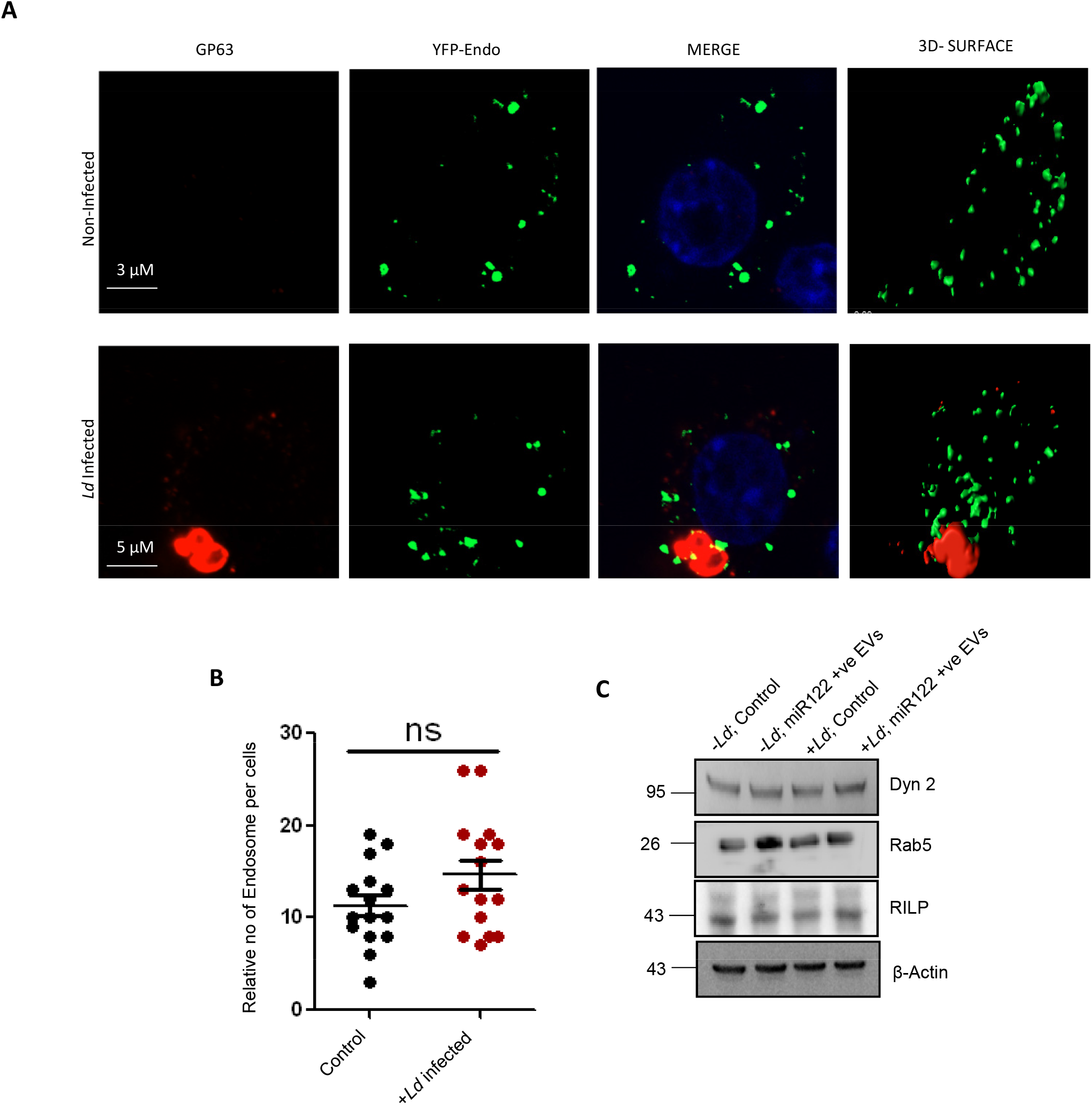
Effect of *Ld* infection on the endosomes of RAW264.7 cells. (A) Effect of *Leishmania* on the endosome number of RAW macrophages. Microscopic images to show the endosomes (YFP-Endo, green) and *Leishmania* parasite (GP63 protein, red). Scale bar 3µm for uninfected and 5µm for infected RAW cells expressing YFP-Endo protein. (B) Graphical representation of the number of endosomes in uninfected or infected macrophages (n=15; p=0.145). (C) Western blots to show the levels of different endocytic pathway component proteins (Dynamin2, Rab5a and RILP) under different conditions of infection and EV-treatment in RAW264.7 cells. Error bars are represented as mean ± S.E.M. P-values were calculated by utilizing two-tailed Student’s t-test. ns: non-significant, *P < 0.05, **P < 0.01, ***P < 0.0001.

**Figure S3.**
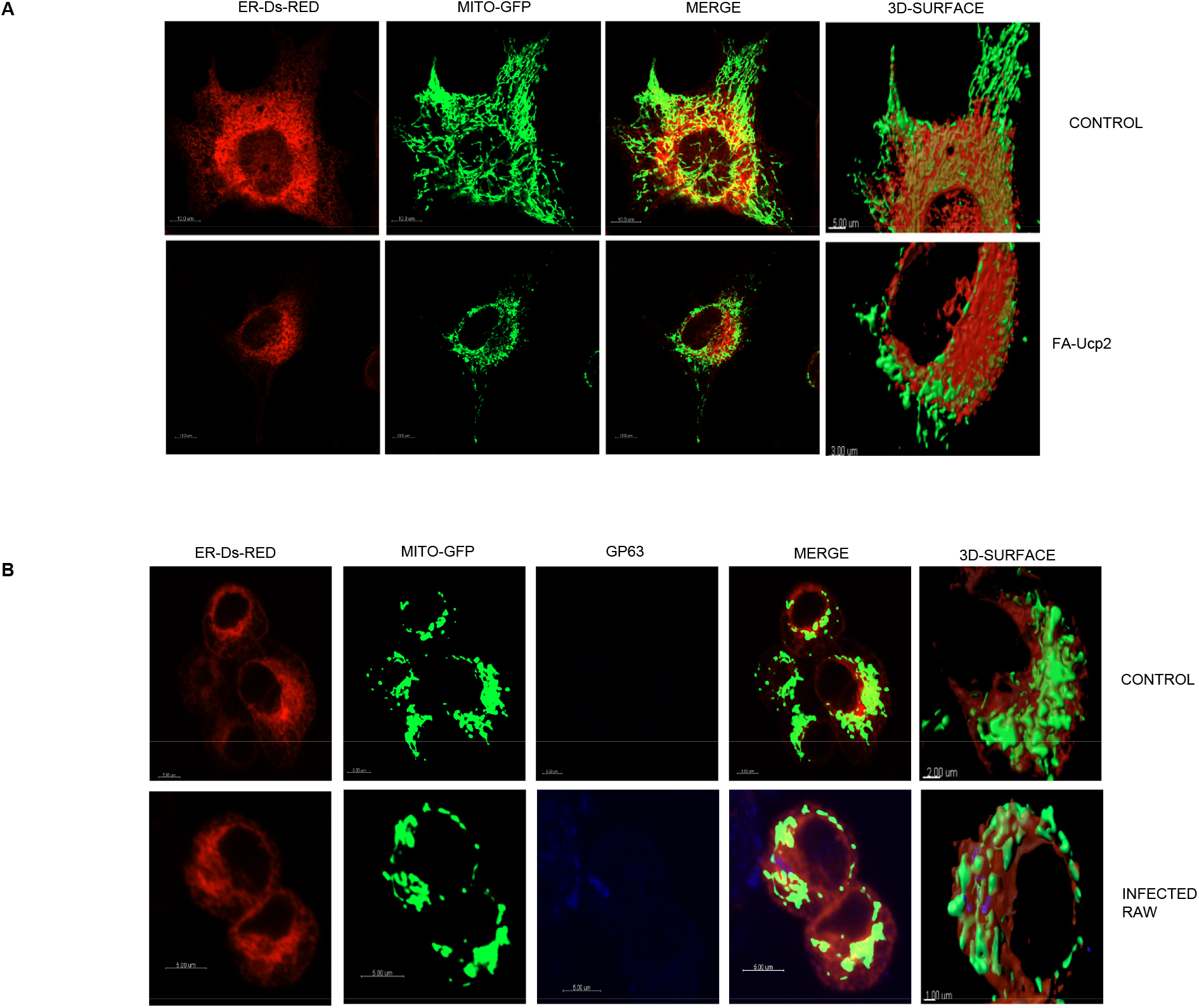
Effect of FA-Ucp2 expression and *Leishmania* infection on ER-mitochondria interaction in HeLa and RAW 264.7 cells. (A) Microscopy of the Endoplasmic Reticulum (red) and mitochondria (green) interaction in Mito-GFP expressing HeLa cells untransfected or transfected with FA-Ucp2. Scale bar 10µm; 5µm for control zoomed surface image and 3µm for FA-Ucp2 expressed zoomed surface image. (C) Microscopy of RAW264.7 cells to show the ER (red) and mitochondria (green) interaction in cells uninfected or infected with *Ld*. The Leishmanial protein GP63 was depicted in blue to show the parasite. Scale bar 5µm; 2µm for control zoomed surface image and 1µm for *Ld* infected zoomed surface image.

**Figure S4.**
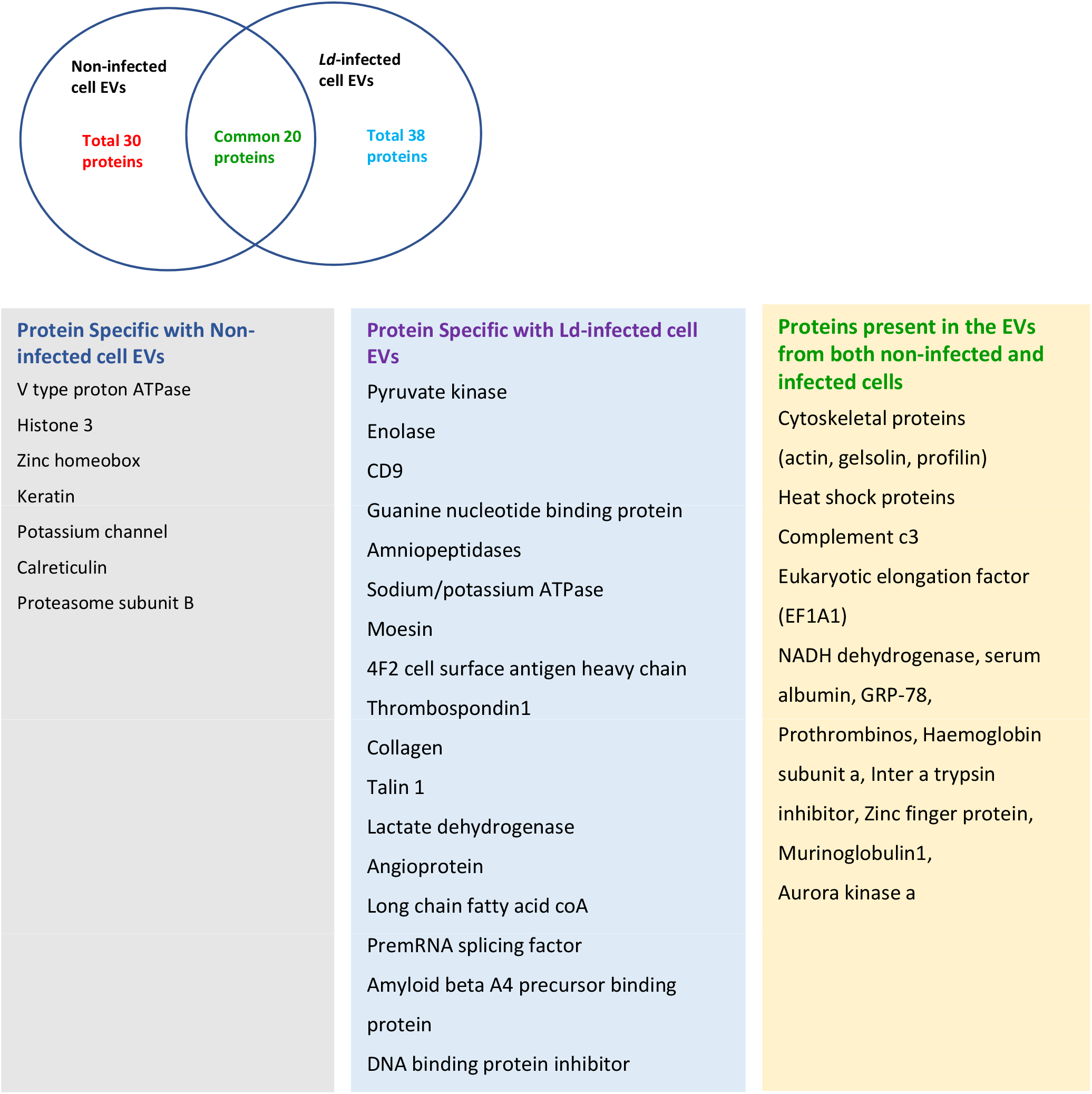
Proteomics profile of control and infected cell derived EVs. EVs were isolated from control and infected cells and a pooled sample of two 90 mm was used for proteomics analysis. Ven diagram showed 20 common proteins of both control and infected cell derived EVS. The list showed the types of proteins found specifically in both types of EVs along with common proteins.

**Figure S5.**
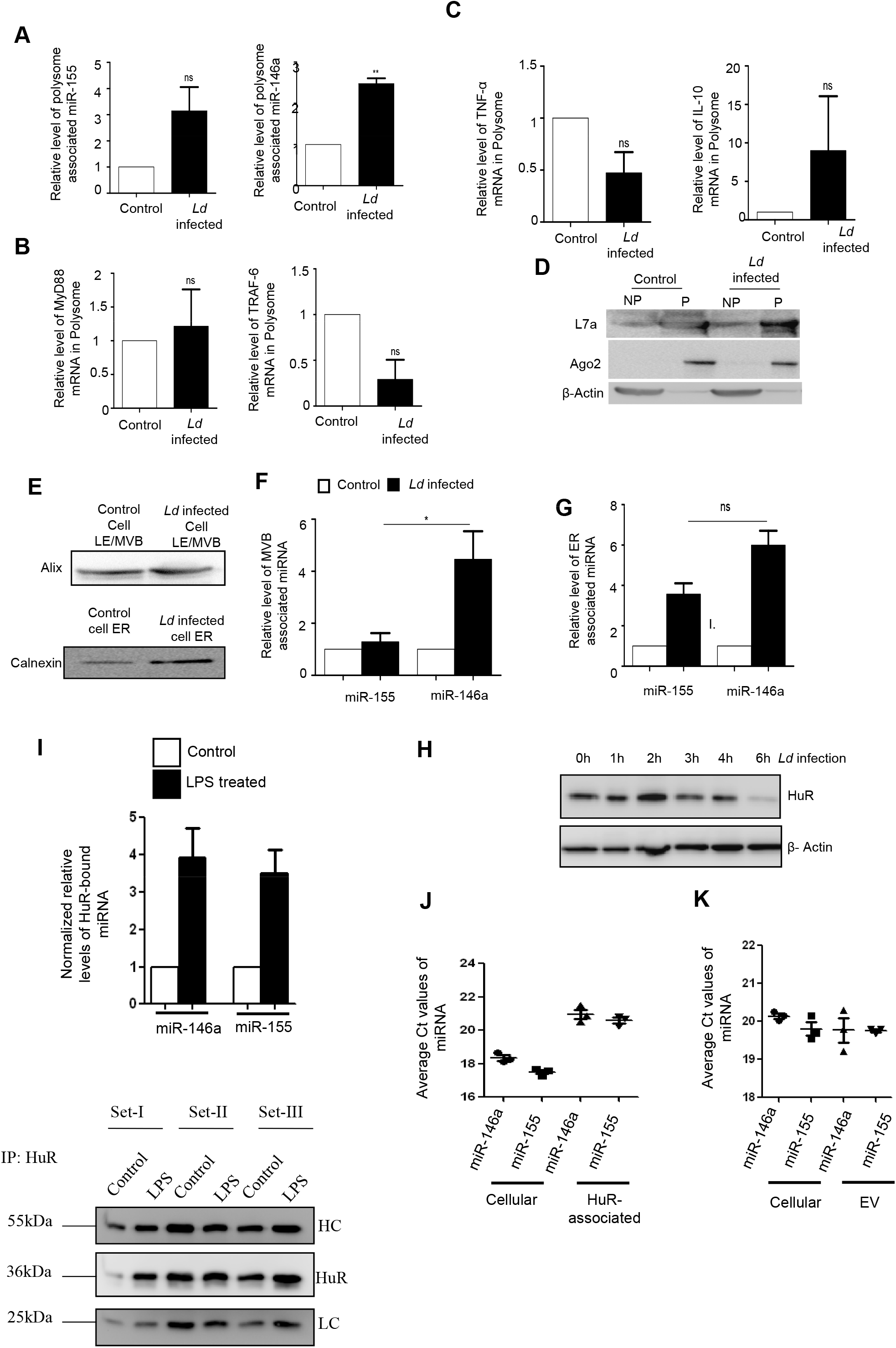
Subcellular sequestration of cytokine mRNAs, microRNAs and their targets in *Ld* infected macrophage cells. (A-D) Polysomal sequestration of cytokine mRNAs, microRNAs and their targets after 24h of *Ld* infection. Relative levels of miR-155 and miR-146a in polysomes were measured by qRT-PCR (A, left panel, p=0.0775, N=5 and right panel, p=0.0077, N=3 respectively). Relative levels of MyD88 and TRAF6 mRNA levels were determined by qRT-PCR in polysome (B, left panel, p=0.7312, N=3 and right panel, p=0.0802, N=3 respectively). Relative cytokine mRNA levels of TNF-α (C, left panel, p=0.0783, N=4), IL-10 (C, right panel, p=0.3740, N=3) were measured by qRT-PCR that are present with polysome in control and infected cells… For mRNA, 18s or GAPDH was used for normalization and for miRNA, polysomal marker L7a was used for normalization (mean+/− S.E.M). Values for uninfected control cells were set as units. Polysomal proteins were determined by western blot (D). (E-G) Subcellular localization of miR-146a and miR-155 in control and infected RAW264.7 cells. Western blot analysis revealed the marker proteins of MVB and ER (E). Relative miR-155 and miR- 146a levels were determined by qRT-PCR of MVB and ER fractions of control and infected cells (F, p=0.0234, N=3, and G, p=0.0514, N=3, unpaired *t*-test). MicroRNA levels were normalized by corresponding marker proteins. Values of control cells were set as units (mean+/−S.E.M). (H) Levels of HuR change with time of infection in *Ld* infected cells. Cell lysates were western blotted for HuR and β-Actin was used as loading control. (I) Increased association of miR-146a and miR-155 with HuR in LPS-treated cells. RAW264.7 cells were treated with LPS and HuR is immunoprecipitated and the amount of miRNAs associated with HuR in control and LPS treated cells are measured by qRT-PCR (upper panel, n=3). The amount of HuR-associated miRNA were normalized against the amount of HuR immunoprecipitated (lower panel) (mean+/−S.E.M, n=3). (J-K) Relative association of miR-146a and miR-155 with HuR and EVs secreted from the HA-HuR expressing RAW264.7 cells. The Ct values of specific miRNAs were plotted for cellular and immunoprecipitated material from cell equivalent amounts of samples (J). The Ct values of the cellular and EV-associated miRNAs in HA-HuR transfected RAW264.7 cells(K) (mean+/−S.E.M, n=3). Amount of HuR immunoprecipitated materials are also shown. In all experimental data, ns: non-significant, *, ** and *** represent *P*-value of <0.05, <0.01 and <0.0001 respectively calculated by student paired *t-*test where not mentioned otherwise.

**Figure S6.**
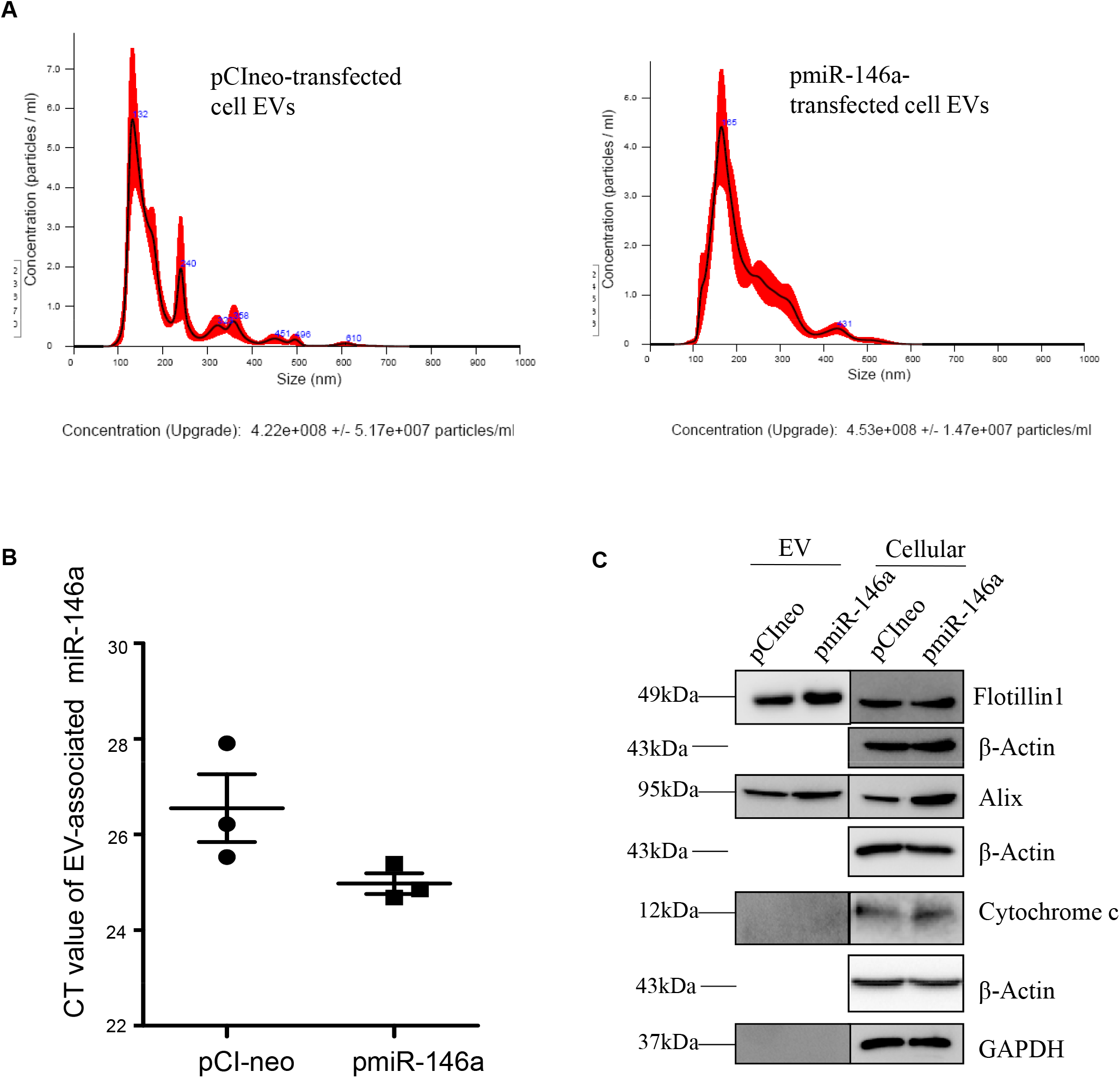
miR-146a does not change EV number or concentration in macrophage cells. (A) Nanoparticle tracking analysis for EVs done for EVs isolated from pCIneo (left panel) or pmiR- 146a (right panel) transfected RAW264.7 cells. The size and concentration of the respective EVs were compared (n=2). PBS was used as blank for nanoparticle tracking analysis. (B) Ct value of EV-associated miR-146a level in pCIneo or pmiR-146a transfected RAW264.7 cells (N.B. – done by Satarupa di) (C) Immunoblotting was done for specific marker proteins which are known to be present and absent in EVs. Both cellular and EV isolated from pCIneo and pmiR-146a transfected RAW264.7 cells were analysed for the respective marker proteins. β-Actin was used as loading control for the cellular samples.

**Figure S7.**
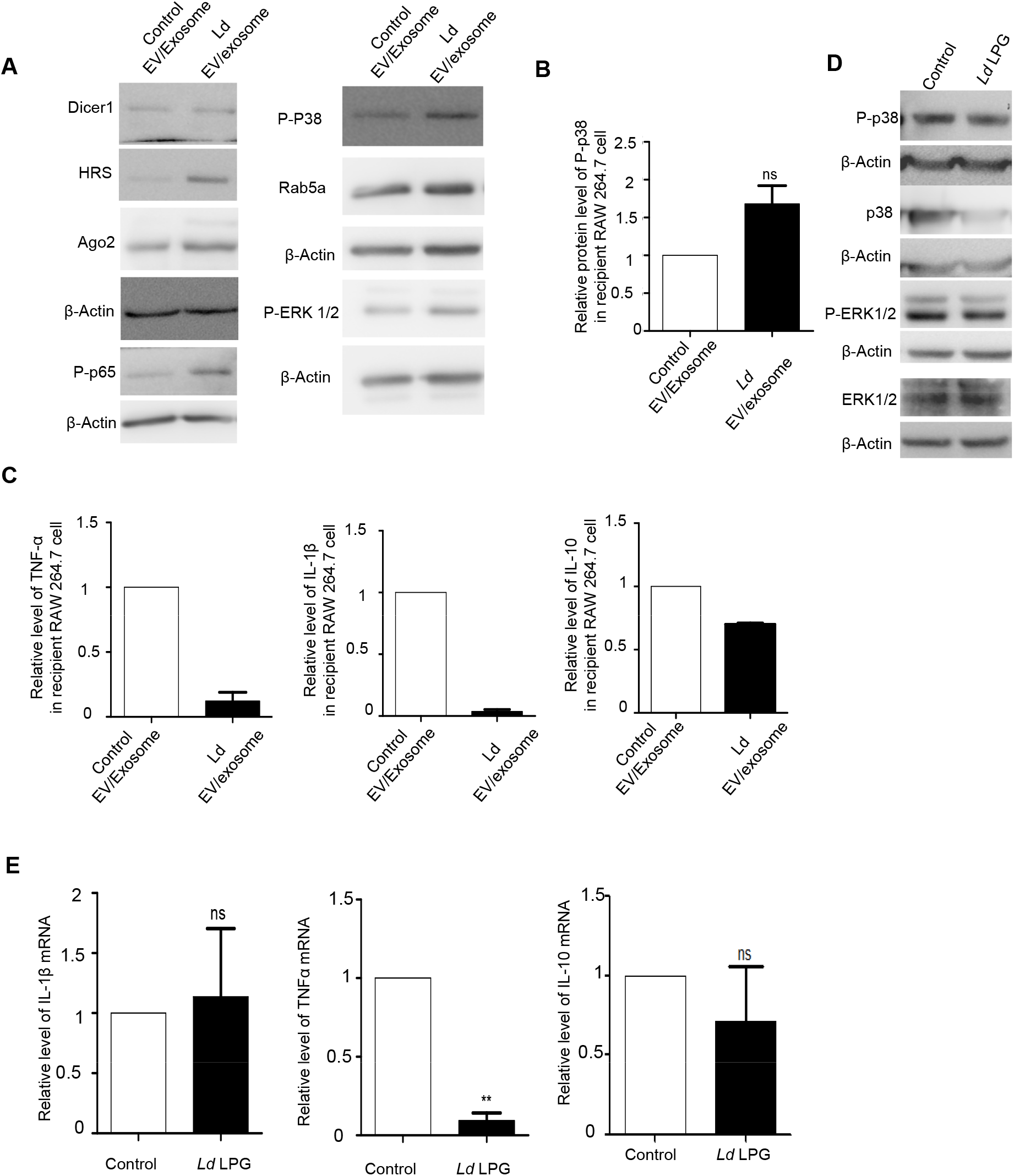
Effect of parasite derived EVs and LPG on cytokine mRNA level and cellular proteins. (A-C) Effect of parasite EVs on cellular proteins and cytokine levels. Protein levels of Dicer1, HRS, Ago2, P-p38, P-ERK1/2, P-p65, Rab5a were determined by western blot of control and *Ld* EV treated macrophages. β-Actin was used as loading control (A). Relative level of P-p38 protein in recipient macrophages treated with control and *Ld* EVs was determined by western blot and band intensities were normalized against actin (B, p=0.1041, N=3). Control EV treated cells were considered as units. Relative cytokine mRNA levels of TNF-α (C, left panel, N=2), IL-1β (C, middle panel, N=2) and IL-10 (C, right panel, N=2) were measured by qRT-PCR in recipient macrophages treated with parasite EVs and medium control EVs for 24h. GAPDH was used as endogenous control and values for control EV treated cells were considered as units (mean+/− S.E.M). (D-E) Effect of *Ld* derived LPG treatment on cellular proteins and cytokine mRNAs. Crude LPG was purified from *Ld* lysate and resuspended in sterile PBS before treatment. RAW264.7 cells were left untreated or treated with LPG at a dose of 1:50 (macrophage: *Ld* cell derived LPG) for 24h and cellular protein levels (P-P38, P-38, P-ERK1/2 and ERK-1/2) were determined by western blots. β- actin was used as loading control (D). Relative mRNA level of IL-1β (E, left panel, p=0.8346, N=3), TNF-α (E, middle panel, p=0.0027, N=3) and IL-10 (E, right panel, p=0.4919, N=3) were determined by qRT-PCR. GAPDH was used as endogenous control and values for untreated cells were considered as units (mean+/− S.E.M). In all experimental data, ns: non-significant, *, ** and *** represent *P*-value of <0.05, <0.01 and <0.0001 respectively calculated by student paired *t-*test where not mentioned otherwise.

**Figure S8.**
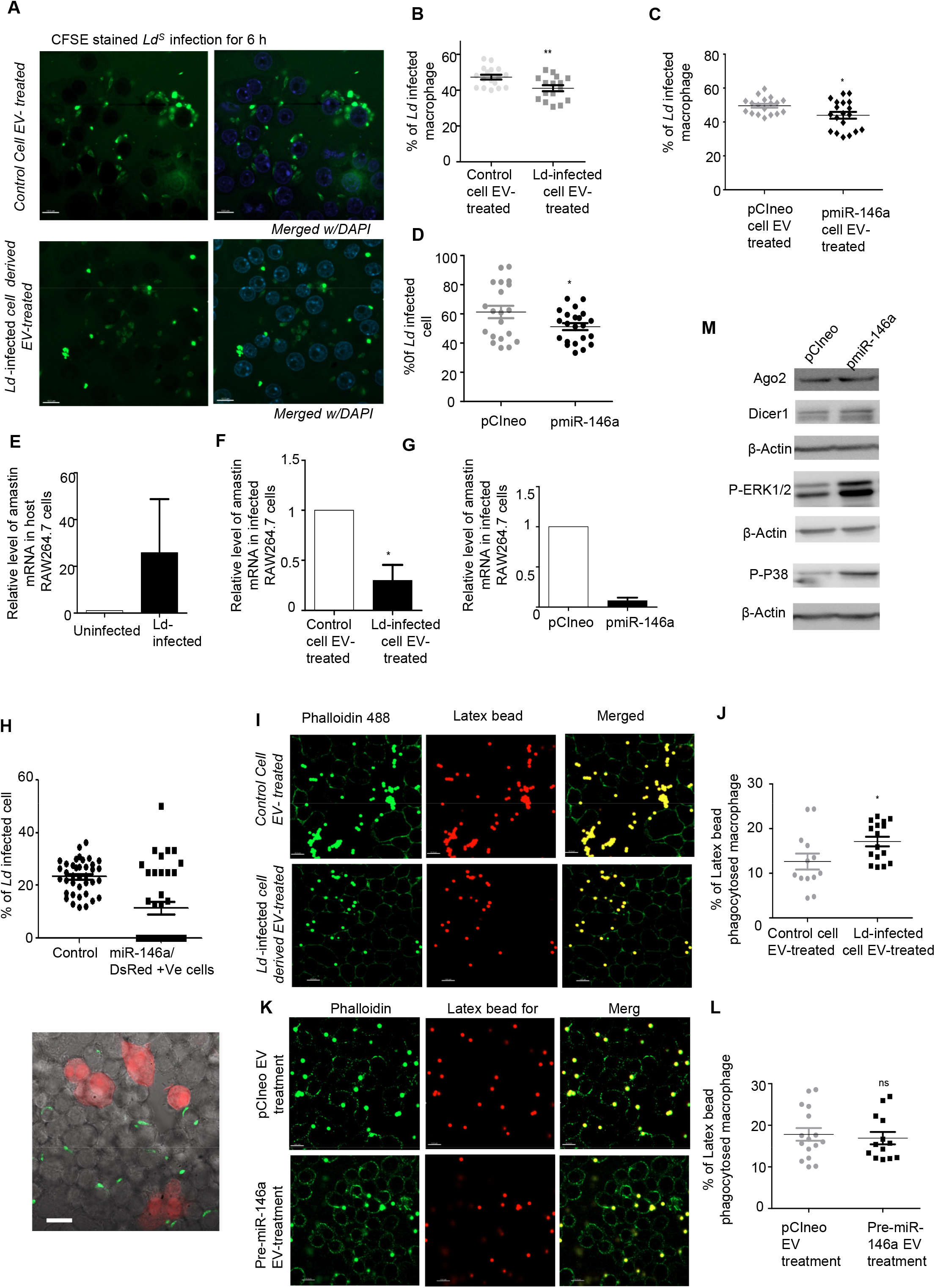
Effect of EV treatment on *Ld* infection. (A-D) Infected cell derived EV treatment decreased *Ld* infection in recipient cells. Parasites were labelled with CFSE dye and infection was given for 6 hours after 24 hours of control and infected cell derived EV treatment (A, upper & lower panel respectively). Parasite infection was determined by counting of CFSE positive structures inside cells. Percentage of infected cells was compared between control cell EV and infected cell EV treated cells (B, p=0.0064, Number of fields, N=16). Parasites were labelled with CFSE dye and infection was given for 6 hours after 24 hour of pCI-Neo and Pre-miR-146a transfected cell derived EV treatment. Percentage of infected cells was compared between pCI-Neo and Pre-miR-146a transfected cell derived EV treated cells (C, p=0.0208, Number of fields, N=18). Parasites were labelled with CFSE dye and infection was given for 24h to pCI-neo and pmiR- 146a transfected cell. Percentage of infected cells was compared between pCI-neo and Pre-miR- 146a transfected cells (D, p=0.0401, Number of fields, N=20). (mean+/− S.E.M) (E-G) RAW264.7 cells were infected with *Ld* for 24hours and infection was measured in terms of amastin mRNA level by qRT-PCR (E, N=2). Relative level of amastin mRNA was determined by qRT-PCR from control and infected cell derived EV treated cells followed by 6h infection (F, p=0.0205, N=4, paired *t*-test). Relative level of amastin mRNA was determined by qRT-PCR from pCI-neo and pmiR-146a transfected cells were followed after 24h of infection (G, N=2). GAPDH was used as endogenous control. Values for uninfected cell or control EV treated cell or pCI-neo transfected cells were considered as units (mean+/− S.E.M). (H) RAW264.7 cells were co-transfected with DsRed and pmiR-146a and was infected with *Ld* for 24h and infection level was visualized at individual cells (lower panel) and quantified (upper panel). Percent of non-miR-146a/DsRed expressing vs non-transfected cells were compared. Scale bar 10µm. (I-L) Latex bead phagocytosis after EV-treatment. Red fluorescent bead (Fluorescent Red, Sigma-Aldrich) was diluted in medium and added to EV-treated RAW264.7 cells at 1:10 (cell:bead) ratio for 6 h. Red fluorescent bead was visualized by a confocal microscope. Phalloidin 488 was used for staining the actin cytoskeleton of cells (I). Percentage of beads phagocytosed in EV treated macrophages was plotted and compared between control cell EV-treated and *Ld* infected cell EV treated cells (J, unpaired *t-*test, p=0.0347, Number of fields, N=13). In panel K visualization of beads phagocytosed in EV treated macrophages was monitored in pCI-neo transfected control cell EV treated and pmiR-146a expressing cell EV treated cells. The quantification has been plotted in panel L (unpaired *t-*test, p=0.6761, Number of fields, N=13). Scale bar=10 µm (mean+/− S.E.M). (M) Cellular protein levels in pmiR-146a and pCI-neo expression conditions were determined by western blot. β-Actin was used as loading control. In all experimental data, ns: non-significant, *, ** and *** represent *P*-value of <0.05, <0.01 and <0.0001 respectively calculated by student unpaired *t-*test where not mentioned otherwise

**Table S1.** Proteomics profile of control and infected cell derived EVs. Detail of the proteins identified.

**Table S2.** Comparative analysis of common proteins present in control as well as infected EVs based on their score of relative abundance and function. Relative score for infected EV is calculated compared to control EV.

| SL. NO | Proteins | EV type | Function | Relative score for infected EV |
| --- | --- | --- | --- | --- |
| 1 | GRP78 | Control & infected | Molecular Chaperon involve in protein folding | 0.76 |
| 2 | Serum Albumin | Control & infected | Carrier protein | 0.96 |
| 3 | Prothrombin | Control & infected | Coagulation factor | 0.8 |
| 4 | Haemoglobin subunit alpha | Control & infected | oxygen carrier | 0.5 |
| 5 | Inter-alpha-trypsin inhibitor heavy chain H3 | Control & infected | Protease inhibitor | 0.9 |
| 6 | Gelsolin | Control & infected | Actin binding protein | 1.13 |
| 7 | Complement C3 | Control & infected | Activator of classical & alternative pathways: Elimination of immunocomplexes | 0.39 |
| 8 | Inter-alpha-trypsin inhibitor heavy chain H2 | Control & infected | Protease inhibitor | 0.51 |
| 9 | Alpha-2-HS-glycoprotein | Control & infected | Carrier protein, role in endocytosis. Brain development, bone tissue formation | 0.61 |
| 10 | Actin, cytoplasmic 1 | Control & infected | Cytoskeletal protein | 0.73 |
| 11 | Profilin-1 | Control & infected | Regulate actin polymerization | 1.66 |
| 12 | AN1-type zinc finger protein 1 | Control & infected | Regulates cytoplasmic stress granule turnover | 1.025 |
| 13 | Protein S100-A6 | Control & infected | Calcium binding proteins | 1.69 |
| 14 | Murinoglobulin-1 | Control & infected | Protease inhibitor | 0.88 |
| 15 | NADH dehydrogenase [ubiquinone] 1 alpha subcomplex assembly factor 4 | Control & infected | Involve in Electron Transport Chain | 0.9 |
| 16 | Adenosylhomocysteinase | Control & infected | enzyme | 1.7 |
| 17 | Alpha-2-macroglobulin-P | Control & infected | antiprotease | 1.19 |
| 18 | Elongation factor 1-alpha 1 | Control & infected | Translation | 6.58 |
| 19 | Aurora kinase A | Control & infected | Regulates cell cycle by regulating mitotic spindles | 1.07 |
| 20 | FtsJ methyltransferase domain-containing protein 1 | Control & infected | Methyltransferase, RNA methylation, Ribosome biogenesis | 1.16 |

**Table S3.** Comparative analysis of proteins present in control and infected EVs based on their score of relative abundance.

| SL No. | Proteins | EV type | Function | score |
| --- | --- | --- | --- | --- |
| 1 | Heat shock-related 70 kDa protein 2 | Control | Molecular chaperon | 100.82 |
| 2 | Heat shock cognate 71 kDa protein | Infected | Protein folding | 126.14 |
| 3 | V-type proton ATPase 16 kDa proteolipid subunit | Control | Protein sorting, receptor mediated endocytosis, synaptic vesicle proton gradient generation | 61.25 |
| 4 | Histone H3.3 | Control | Genome integrity | 32.24 |
| 5 | Zinc finger homeobox protein 3 | control | Transcriptional regulator | 27.82 |
| 6 | Keratin, type II cytoskeletal 74 | control | Hair formation | 27.75 |
| 7 | Potassium voltage-gated channel subfamily KQT member 5 | control | Ion transport | 24.93 |
| 8 | Protein unc-80 homolog | control | Cation channel activity | 24.93 |
| 9 | Calreticulin | control | Calcium binding | 24.65 |
| 10 | MAP kinase-activating death domain protein | control | Guanyl nucleotide exchange factor | 23.84 |
| 11 | Proteasome subunit beta type-10 | control | Peptidase activity | 21.83 |
| 12 | Pyruvate kinase isozymes M1/M2 | infected | glycolysis | 282.66 |
| 13 | Alpha-enolase | infected | glycolysis | 198.56 |
| 14 | CD9 antigen | infected | Tetraspanin protein | 179.48 |
| 15 | Guanine nucleotide-binding protein G(I)/G(S)/G(T) subunit beta-2 | infected | G-protein formation | 89.33 |
| 16 | Aminopeptidase N | infected | Peptidase activity, role in antigen presentation, angiogenesis | 43.62 |
| 17 | Sodium/potassium-transporting ATPase subunit alpha-1 | infected | Ion transport channel formation | 43.47 |
| 18 | Moesin | infected | T cell & B cell homeostasis, self tolerance | 39.66 |
| 19 | 4F2 cell-surface antigen heavy chain | infected | Amino acid transporter | 36.50 |
| 20 | Thrombospondin-1 | infected | Adhesive glycoprotein | 35.67 |
| 21 | Collagen alpha-1(VI) chain | infected | Tissue integrity | 34.38 |
| 22 | Talin-1 | infected | Linking integrin with actin | 32.77 |
| 23 | L-lactate dehydrogenase C chain | infected | Pyruvate metabolism | 25.39 |
| 24 | Angiopoietin-related protein 4 | infected | Angiogenesis, lipid metabolism | 24.90 |
| 25 | Pre-mRNA-splicing factor ISY1 homolog | infected | mRNA splicing | 24.65 |
| 26 | Long-chain-fatty-acid--CoA ligase 1 | infected | Lipid metabolism | 23.11 |
| 27 | Amyloid beta A4 precursor protein-binding family A member 3 | infected | Protein transport | 22.14 |
| 28 | DNA-binding protein inhibitor ID-4 | infected | Transcription regulation | 20.94 |

**Supplementary Table S4.** Details of miRNA assays used for Taqman based quantification.

| miRNA | Assay ID |
| --- | --- |
| mouse miR-155 | 002571 |
| human miR-146a | 000468 |
| human miR-21 | 000397 |
| human miR-16 | 000391 |
| human miR-125b | 000449 |
| human miR-122 | 000445 |
| U6 | 001973 |

**Supplementary Table S5.**
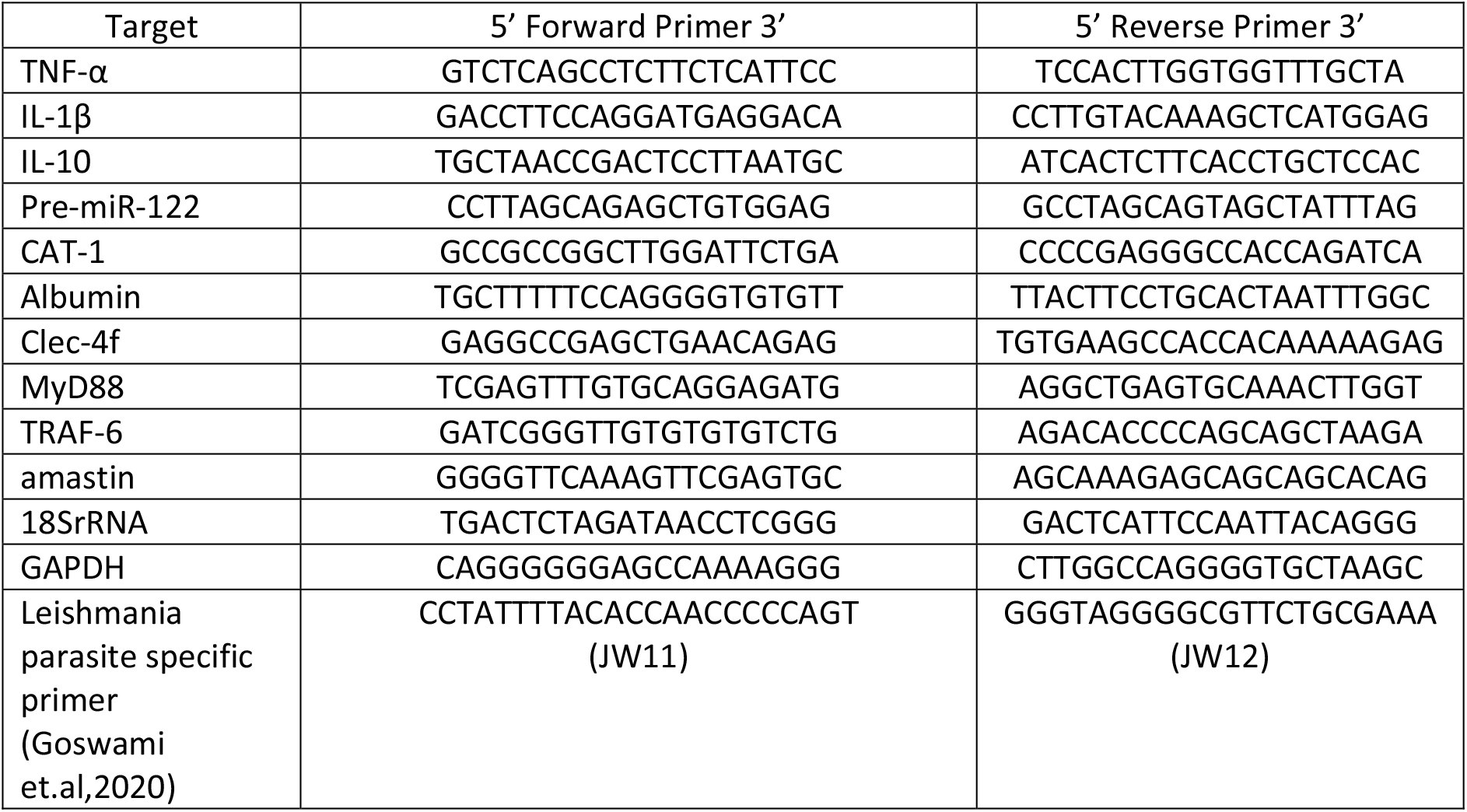
Details of mRNA primers used for SYBR-Green based quantification.

**Supplementary Table S6.**
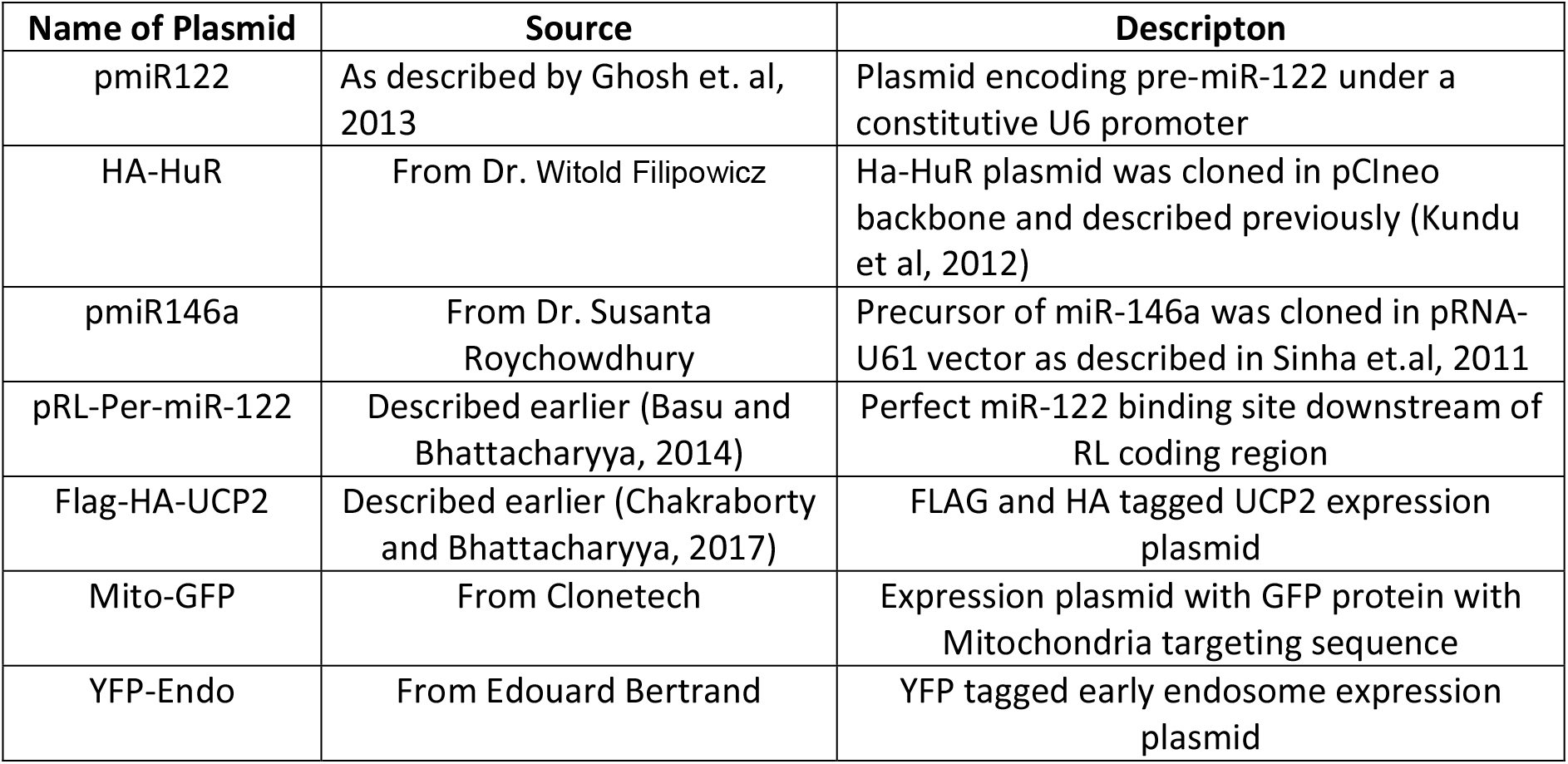
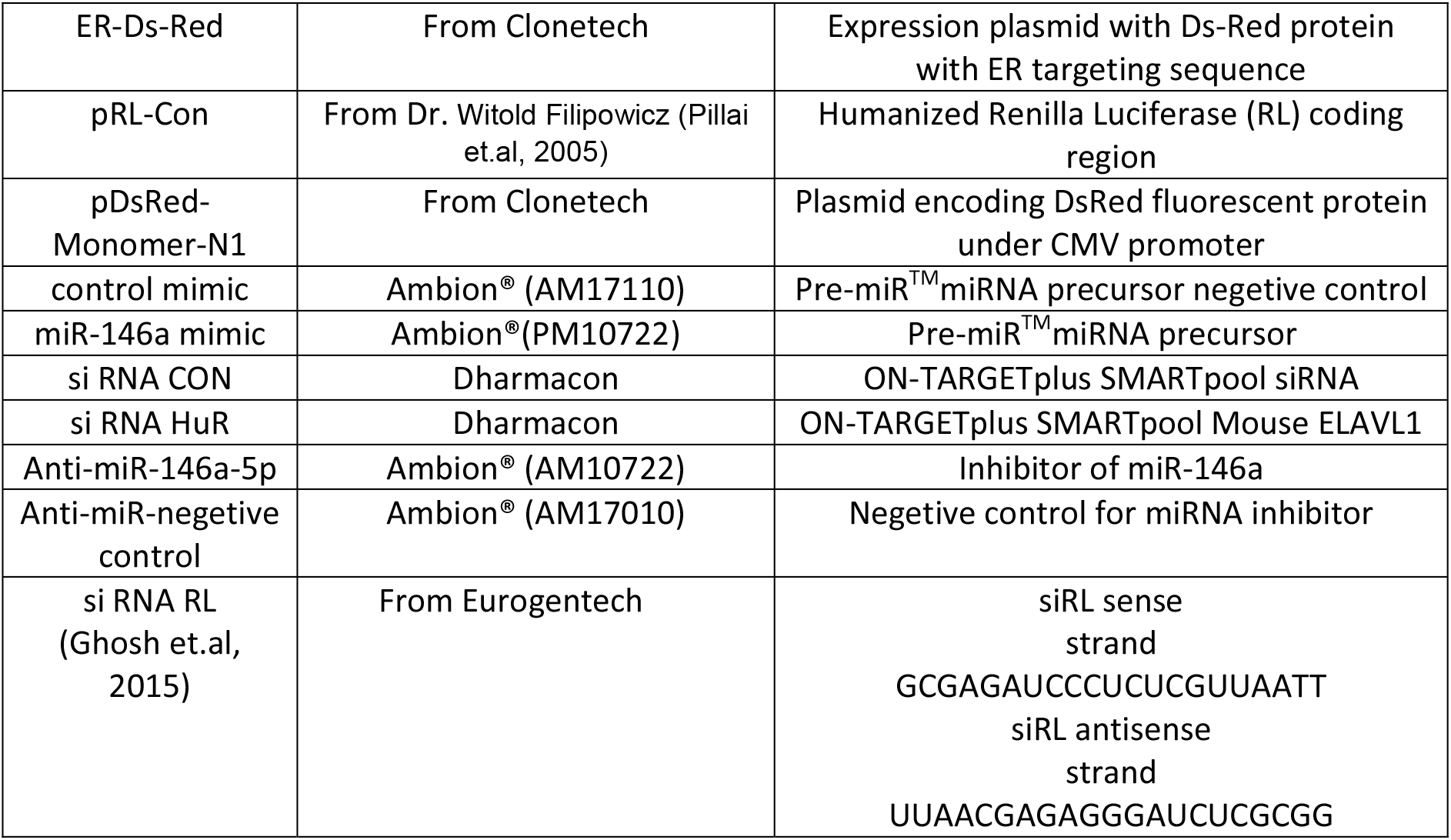
List of Plasmids, siRNAs, miRNA mimics and miRNA inhibitor.

**Supplementary Table S7.** Antibodies used for Western Blot.

| Antibody name | Raised in | Dilution used | Source |
| --- | --- | --- | --- |
| Dicer1 | Rabbit | 1:5000 | Bethyl |
| HRS | Rabbit | 1:1000 | Bethyl |
| RAB27a | Rabbit | 1:1000 | Cell Signaling |
| Flotilin 1 | Rabbit | 1:1000 | Cell Signaling |
| Alix | Mouse | 1:200 | Santa Cruz |
| Cytochrome C | Rabbit | 1:1000 | Cell Signaling |
| HuR | Mouse | 1:1000 | Santa Cruz |
| P-p38 | Rabbit | 1:1000 | Cell Signaling |
| P-ERK1/2 | Rabbit | 1:1000 | Cell Signaling |
| HA | Rat Monoclonal | 1:1000 | Roche |
| ERK1/2 | Rabbit | 1:1000 | Cell Signaling |
| P-38 | Rabbit | 1:1000 | Cell Signaling |
| L7a | Rabbit | 1:1000 | Cell Signaling |
| Ago2 | Mouse | 1:1000 | Abnova |
| Calnexin | Rabbit | 1:8000 | Bethyl |
| Rab5 | Rabbit | 1:1000 | Cell Signaling |
| GAPDH | mouse | 1:6000 | Sigma-Aldrich |
| CD63 | Mouse | 1:1000 | BD Pharmingen |
| β-actin | Mouse monoclonal (HRP conjugated) | 1:10000 | Sigma Aldrich |
| UCP2 | Goat | 1:5000 | Novus |
| Rab7 | Rabbit | 1:1000 | Cell Signaling |
| Dynamin 2 | Rabbit | 1:1000 | Cell Signaling |
| RILP | Goat | 1:1000 | Santa Cruz |
| GP63 | Mouse | 1:1000 | LS Biosciences |

